# Bidirectional control of a metabolic transition by the GID ubiquitin ligase

**DOI:** 10.1101/2025.07.15.664932

**Authors:** Ka-Yiu Edwin Kong, Christian Ochs, Cécile Debarnot, Matti Myllykoski, Denís Arribas Blanco, Susmitha Shankar, Vivien A.C. Schoonenberg, Julia Kończak, Julia K. Varga, Ora Schueler-Furman, Katja Luck, Falk Butter, Jia-Xuan Chen, Lukas S. Stelzl, Thomas Arnesen, Anton Khmelinskii

## Abstract

The GID/CTLH ubiquitin ligase is a multisubunit E3 conserved across eukaryotes. GID/CTLH has been implicated in a variety of processes, including metabolic regulation, cell proliferation, embryonic development and cell differentiation. However, our understanding of substrate recognition by GID/CTLH remains incomplete. Here, we characterize the regulation, specificity and functions of the putative substrate receptor Gid11 in budding yeast. We find that upon its expression during the switch of carbon source from glucose to ethanol, Gid11 associates with the core GID complex, likely forming a large ring assembly. Through systematic mutagenesis, we identify a negatively charged pocket on Gid11 that binds substrates carrying N-terminal degrons starting with threonine and define the Thr/N-degron motif. Such Thr/N-degrons are exposed upon co-translational removal of the initiator methionine, while escaping N-terminal acetylation. Finally, we show that during the switch to ethanol, GID^Gid11^ targets for degradation factors linked to glycolysis, opposing the established role of GID in the degradation of gluconeogenic enzymes during the reverse switch from ethanol to glucose. Taken together, our results establish Gid11 as a bona fide GID substrate receptor and highlight GID as a regulator of metabolic transitions.

## Introduction

Selective protein degradation is involved in most cellular processes and contributes to proteome homeostasis through the removal of unnecessary or abnormal proteins (Balch et al., 2008; Balchin et al., 2016). The ubiquitin-proteasome system (UPS) plays a key role in selective protein degradation, whereby ubiquitin-protein ligases (ubiquitin ligases or E3s hereafter) mark proteins with ubiquitin for degradation by the 26S proteasome (Zheng and Shabek, 2017). Ubiquitination substrates are typically recognized by E3 substrate receptor subunits via short linear motifs known as degradation signals or degrons. Remarkably, degrons can be as short as one or two amino acids, especially in the case of degrons found at protein termini (N- and C-degrons at protein N- and C-termini, respectively) (Sherpa et al., 2022; Timms and Koren, 2020; Varshavsky, 2019).

Numerous ubiquitin ligases, including cullin RING ubiquitin ligases (CRLs) and the anaphase promoting complex/cyclosome (APC/C), are modular multisubunit assemblies that recruit substrates using interchangeable substrate receptors, thus expanding their substrate repertoire (Alfieri et al., 2017; Harper and Schulman, 2021; Wang et al., 2020). As these receptors can compete for the same binding site on the E3, assembly of substrate-specific E3 complexes can be dynamic and regulated through mechanisms that include conditional expression of a given substrate receptor. The GID (glucose-induced degradation deficient) ubiquitin ligase has recently emerged as an example of E3s employing interchangeable substrate receptors (Liu and Pfirrmann, 2019; Maitland et al., 2022; Sherpa et al., 2022). The GID complex is conserved from yeast to humans, where it is known as the CTLH (C-terminal to LisH) complex. Both GID and CTLH subunits can assemble into multiple distinct large complexes that appear to recognize and ubiquitinate different substrates (Maitland et al., 2022). Thus, GID/CTLH ubiquitin ligases have been implicated in a variety of processes, including metabolic regulation, cell proliferation, embryonic development and cell differentiation. Furthermore, mutations in CTLH subunits have been linked to numerous pathologies, including cancer and neurodevelopmental disorders (Huffman et al., 2019; Maitland et al., 2022).

In the budding yeast *Saccharomyces cerevisiae*, there are at least three functionally distinct GID assemblies. The core catalytic unit comprises a RING module consisting of the RING and RING-like subunits Gid2 and Gid9, and a substrate receptor module consisting of the Gid1 and Gid8 scaffold subunits and the adaptor subunit Gid5, which recruits the interchangeable substrate receptors Gid4 and Gid10 (Hämmerle et al., 1998; Menssen et al., 2012; Qiao et al., 2020; Regelmann et al., 2003; Santt et al., 2008). The GID^Gid4^ core complex targets for degradation three gluconeogenic enzymes (the malate dehydrogenase Mdh2, the phosphoenolpyruvate carboxykinase Pck1 and the isocitrate lyase Icl1) during the switch of carbon source from ethanol to glucose. Gid4 recognizes these substrates via their N-terminal degrons starting with a proline residue (Pro/N-degrons) (Chen et al., 2017; Hämmerle et al., 1998). Similar to Gid4, the Gid10 receptor is a beta-barrel protein and has a similar specificity towards Pro/N-degrons (Melnykov et al., 2019; Qiao et al., 2020). Importantly, expression of Gid4 and Gid10 is conditional. Whereas Gid4 is upregulated during the switch of carbon source from ethanol to glucose (Menssen et al., 2018; Santt et al., 2008), Gid10 expression is increased upon heat shock, osmotic stress or starvation (Melnykov et al., 2019; Qiao et al., 2020), highlighting the conditional nature of the GID^Gid4^ and GID^Gid10^ assemblies.

With the help of the Gid7 subunit (or its human orthologs WDR26 and MKLN1), two core complexes can form a large ring-like assembly, named the chelator complex (Sherpa et al., 2021). Ubiquitination and subsequent degradation of only one of the Gid4 substrates, the Fbp1 fructose-1,6-bisphosphatase, requires such a large assembly. The two Gid4 receptors in the ring complex help orient Fbp1 tetramers for ubiquitination of specific lysine residues. In cells lacking Gid7, Fbp1 is stable whereas Gid4-dependent degradation of Mdh2, Pck1 and Icl1 remains unaffected during the switch of carbon source from ethanol to glucose (Sherpa et al., 2021). Besides this scaffold function, Gid7 orthologs in other species (including flies and humans) can function as substrate receptors (Briney et al., 2025; Gottemukkala et al., 2024; Maitland et al., 2022; Mohamed et al., 2021). Interestingly, human cells can use the pseudo-substrate YPEL5 (Moh1 in yeast) to block the substrate binding site of WDR26 and in this way inhibit degradation of a GID substrate, the nicotinamide/nicotinic-acid-mononucleotide-adenylyltransferase NMNAT1 (Gottemukkala et al., 2024). It is unclear whether yeast Gid7 and Moh1 function similarly. Finally, three chelator complexes can combine into a large assembly with the Gid12 subunit. While the functions of this assembly are not entirely clear, Gid12 inhibits GID^Gid4^-dependent protein degradation (Qiao et al., 2022).

Previously, we uncovered Gid11 as a putative GID substrate receptor in budding yeast and identified 13 proteins with Gid11-dependent turnover (Kong et al., 2021). In contrast to Gid4 and Gid10 substrates, none of the 13 potential Gid11 substrates have an N-terminal proline. One of them, the Cpa1 subunit of the carbamoyl phosphate synthetase, has a serine residue after the initiator methionine. Mutation of this N-terminal serine had no impact on Gid11-dependent turnover of Cpa1. In the other 12 potential Gid11 substrates, the initiator methionine is followed by a threonine residue. Mutating this N-terminal threonine to alanine or glycine in three potential substrates, the putative phosphoglycerate mutase Gpm3, the nucleotidase Phm8 and the phosphatase of unknown function Yor283w, resulted in completely stable proteins. Importantly, Gid4, Gid10 and Gid11 share a C-terminal motif for interaction with Gid5 (Kong et al., 2021; Qiao et al., 2020). These observations led us to propose that Gid11 is a GID substrate receptor recognizing proteins with threonine N-degrons (Thr/N-degrons hereafter) (Kong et al., 2021). Here we define the composition, specificity and functions of the GID^Gid11^ complex.

## Results

### Regulation of Gid11

To follow expression of the Gid11 protein, we tagged *GID11* at its endogenous chromosomal locus with the hemagglutinin (HA) epitope. We then used fluorescence measurements of colonies expressing Gid11 substrates tagged with the mCherry-sfGFP fluorescent timer (tFT) to assess the functionality of tagged Gid11. The tFT is a reporter of protein turnover, whereby tFT-tagged proteins with fast turnover exhibit low mCherry/sfGFP ratios and vice-versa (Khmelinskii et al., 2012). Whereas C-terminally tagged Gid11-HA was not functional, N-terminally tagged HA-Gid11 supported turnover of Gpm3, Phm8 and Yor283w in the tFT colony assay to the same extent as untagged Gid11 (Fig. S1a).

HA-Gid11 was upregulated in conditions such as carbon or nitrogen starvation and osmotic stress (Fig. S1b), in agreement with our previous observations (Kong et al., 2021). From here on, we focused on the switch of carbon source from glucose to ethanol (glucose-to-ethanol switch hereafter), where Gid11 exhibited the strongest upregulation. During this transition, Gid11 levels increased progressively for approximately 6 h and declined afterwards (Fig. 1a). Accumulation of Gid11 protein was preceded by accumulation of *GID11* mRNA (Fig. 1b), suggesting transcriptional control of *GID11*. Consistent with this notion, we observed increased activity of the *GID11* promoter 1h after the glucose-to-ethanol switch using activity of the NanoLuc luciferase expressed from the *GID11* promoter (*GID11pr*) as a readout (Fig. 1c). For comparison, the activity of the *TDH3* promoter (*TDH3pr*) was not affected during this transition.

**Figure 1.**
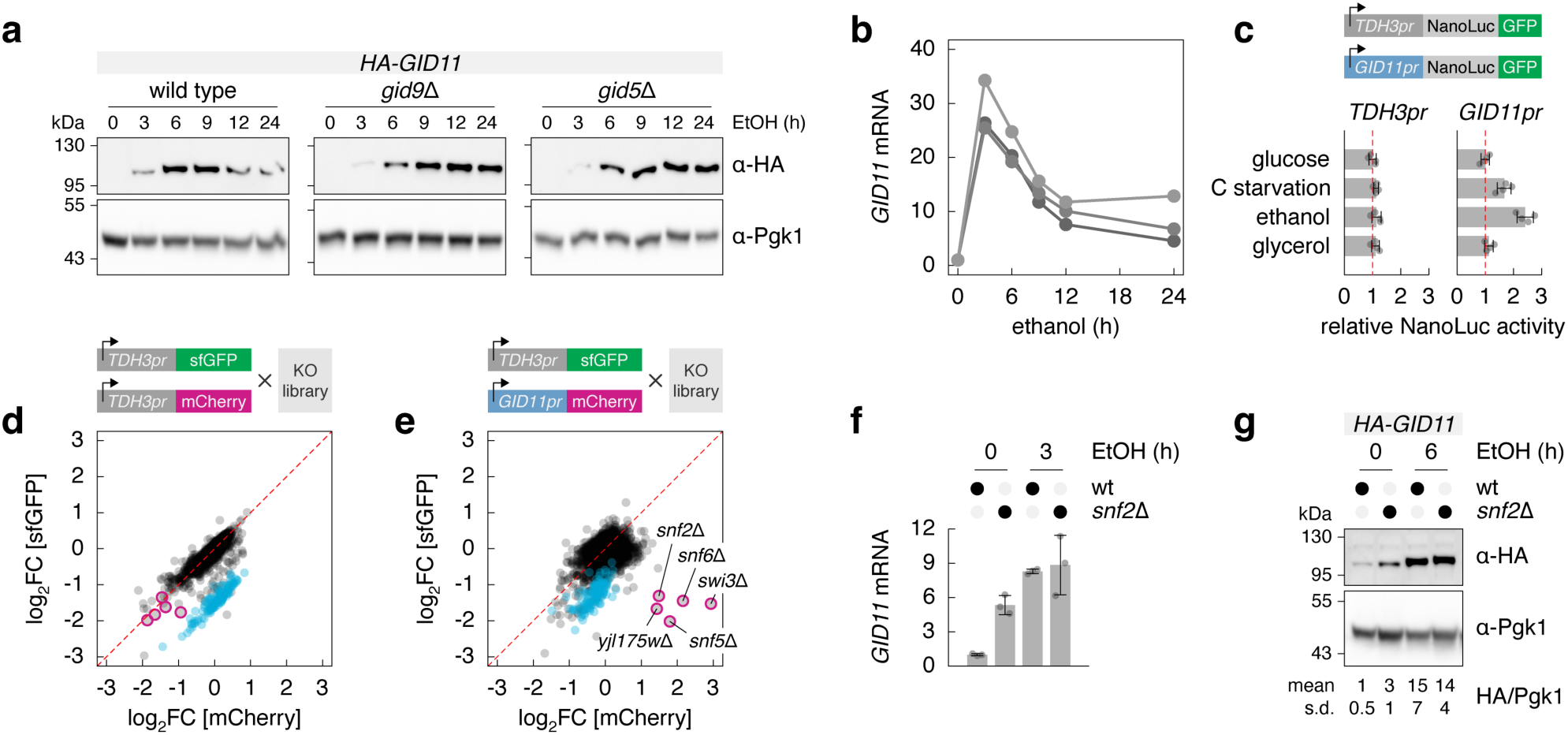
Multiple factors regulate conditional expression of Gid11. **a** – Gid11 protein levels during the switch of carbon source from glucose to ethanol. Whole-cell extracts were separated by SDS-PAGE, followed by immunoblotting with antibodies against the HA tag and Pgk1 as the loading control. **b** – Changes in *GID11* mRNA levels during the switch of carbon source from glucose to ethanol. Quantification of *GID11* mRNA relative to *ALG9* mRNA with RT-qPCR, normalized to time point zero, of three biological replicates. **c** – Activation of the *GID11* promoter by ethanol. Quantification of luciferase activity in strains carrying the indicated constructs and grown on glucose as carbon source, undergoing carbon (C) starvation for 1 h or 1 h after the switch of carbon source to ethanol or glycerol (n = 4 biological replicates). **d, e** – Genome-wide screen for modulators of the *GID11* promoter activity with the yeast non-essential gene knockout library (KO library). Fold changes (FC, log_2_ scale) in fluorescence between each gene knockout and the mean of the corresponding screen plate (n = 4 technical replicates). Mutants with a significant effect in the control screen (**d**, log_2_FC (mCherry/sfGFP) > 1 at 5% FDR) are marked in cyan. Mutants affecting the activity of the *GID11* promoter (**e**) are highlighted with magenta outlines. **f, g** – Quantification of *GID11* mRNA with RT-qPCR (**f**) and HA-Gid11 protein levels by immunoblotting of whole cell extracts (**g**). Measurements normalized to the wild type at time point zero (mean ± s.d., n = 3 biological replicates).

We thus performed a genome-wide screen to identify regulators of *GID11* transcription. We introduced pairs of promoter activity reporters, *TDH3pr-sfGFP* and *GID11pr-mCherry* or *TDH3pr-mCherry*, into the yeast knockout library carrying deletion alleles of non-essential genes (Giaever et al., 2002; Winzeler et al., 1999). Fluorescence measurements of the resulting colony arrays identified subunits of the SWI/SNF chromatin remodeling complex as negative regulators of the *GID11pr* activity (Fig. 1d, e, Supplementary Data 1). Consistent with the screen results, strains lacking the Snf2, Snf5, Snf6 or Swi3 subunits of the SWI/SNF complex exhibited increased mCherry/sfGFP ratios when mCherry was expressed from the *GID11* but not from the *TDH3* promoter (Fig. S1c). To understand when the SWI/SNF complex influences *GID11* expression, we analyzed *GID11* mRNA and protein levels during the glucose-to-ethanol switch. *snf2*Δ strains exhibited increased *GID11* mRNA and protein levels when grown on glucose (5.3±0.8 and 3±1 fold change compared to wild type, respectively; mean ± s.d., n = 3 technical replicates) but not after the glucose-to-ethanol switch (Fig. 1f, g), arguing for transcriptional repression of *GID11* in glucose.

Upregulation of Gid11 was transient, with protein levels peaking 6h after the glucose-to-ethanol switch. Subsequent Gid11 downregulation was lost in strains lacking the RING subunit Gid9 or the adaptor subunit Gid5 (Fig. 1a), suggesting that activity of the GID complex and recruitment of Gid11 to the GID complex are required for Gid11 turnover. This behavior is similar to the other substrate receptors Gid4 and Gid10 (Langlois et al., 2022; Menssen et al., 2018). We conclude that conditional expression of Gid11 is controlled at least in part via transcriptional repression by the SWI/SNF complex in glucose medium and GID-dependent turnover in ethanol medium.

### Composition and function of the GID^Gid11^ complex

We sought to define the composition of the GID^Gid11^ complex. Mass spectrometry analysis of Gid1-HA immunoprecipitates from cells grown on glucose compared to an untagged control strain identified all scaffold GID subunits (Gid1, Gid5 and Gid8), the RING subunits Gid2 and Gid9, the oligomerization subunit Gid7, the Gid4 substrate receptor and Gid12, a steric inhibitor of GID^Gid4^ complexes (Qiao et al., 2022) (Fig. 2a, Supplementary Data 2). Gid11 was identified in Gid1-HA immunoprecipitates only from cells grown on ethanol (Fig. 2b). Interestingly, both Gid4 and Gid11 co-immunoprecipitated with Gid1-HA under this condition.

**Figure 2.**
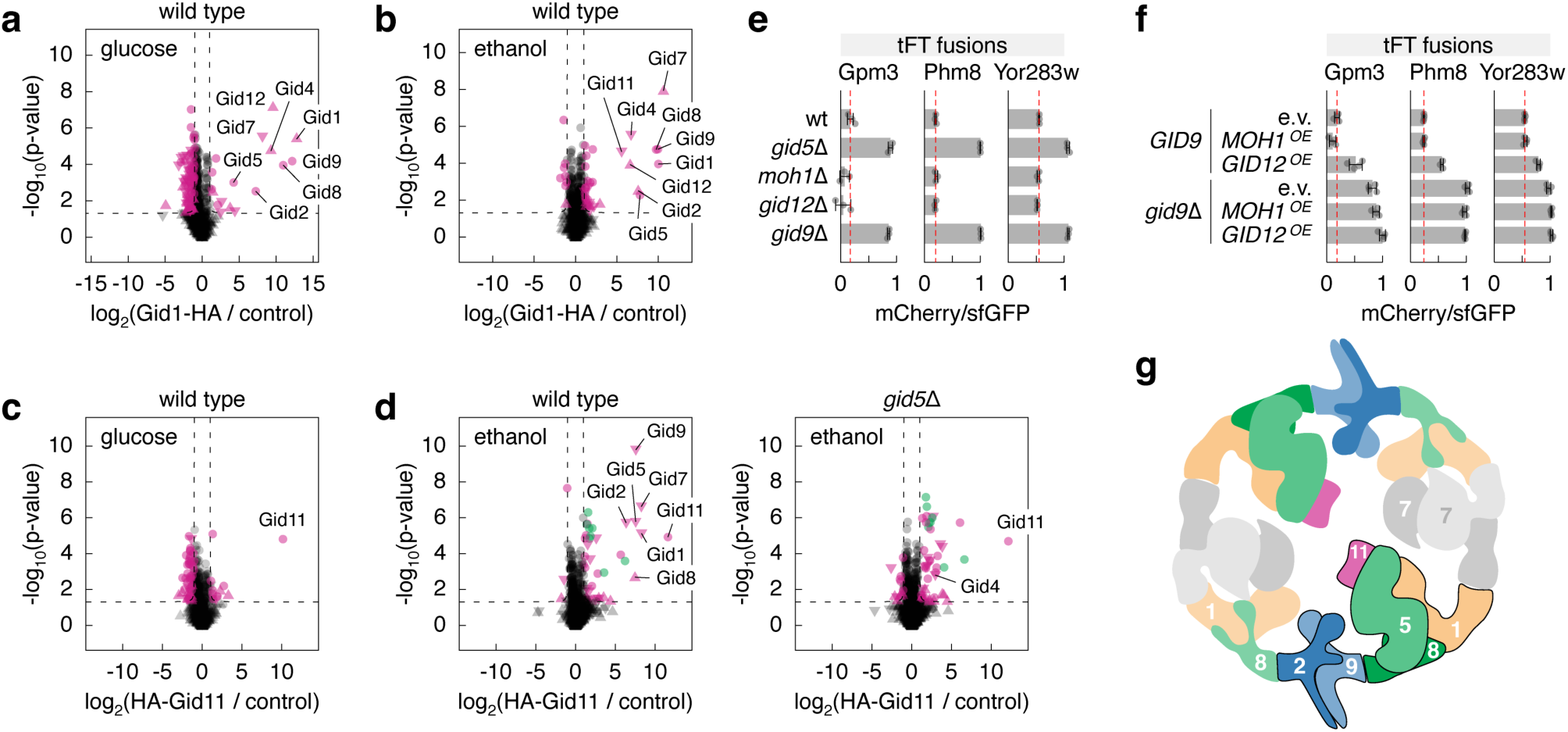
Conditional assembly of the GID^Gid11^ complex. **a-d** – Mass spectrometry analysis of proteins co-immunoprecipitated with Gid1-HA (**a**, **b**) or HA-Gid11 (**c**, **d**). Strains grown on glucose as carbon source (**a**, **c**) or 6 h after switching to ethanol (**b**, **d**). Volcano plots of differences in mean signal intensities between the HA-Gid11 immunoprecipitation and an untagged control (n = 4 technical replicates). Significantly enriched or deleted proteins are marked in magenta (absolute(log_2_(sample/control)) > 1 and p-value < 0.05), subunits of the CCT chaperonin are highlighted in green. **e, f** – mCherry/sfGFP ratios of colonies expressing tFT-tagged proteins (mean ± s.d., n = 4 biological replicates; wt, wild type). OE, overexpression. **g** – Cartoon of the putative GID^Gid11^ ring complex, by similarity with the GID^Gid4^ ring complex (Sherpa et al., 2021).

To determine whether these immunoprecipitates represent multiple assemblies with different substrate receptors, we analyzed the interactome of HA-Gid11. Gid7, the scaffold and RING subunits, but not Gid4, were enriched in HA-Gid11 immunoprecipitates from cells grown on ethanol but not glucose (Fig. 2c, d, Supplementary Data 2). The association of HA-Gid11 with the other GID subunits depended on Gid5 (Fig. 2d). Gid12 and Moh1, the ortholog of human YPEL5, which binds the human Gid7 ortholog WDR26 (Gottemukkala et al., 2024; Lampert et al., 2018; Sherpa et al., 2021), were not enriched in HA-Gid11 immunoprecipitates from wild type cells. Consistent with this, deletion of *MOH1* or *GID12*, or *MOH1* overexpression had no impact on turnover of Gid11 substrates in the tFT colony assay (Fig. 2e, f). Interestingly, overexpression of *GID12* stabilized all tested Gid11 substrates (Fig. 2f), raising the possibility that Gid12 could function as a steric inhibitor of GID^Gid11^ complexes, akin to its ability to inhibit GID^Gid4^ complexes (Qiao et al., 2022). Finally, subunits of the TRiC/CCT chaperonin complex were among HA-Gid11 interactors (Fig. 2d). It is possible that TRiC/CCT contributes to folding of Gid11. Based on these observations and the previously described structure of the GID^Gid4^ chelator complex (Sherpa et al., 2021), we hypothesize that GID^Gid11^ forms an analogous assembly, with two substrate receptor (Gid1, Gid5, Gid8, Gid11) and two RING (Gid2, Gid9) modules dimerized by Gid7, with Gid11 replacing Gid4 (Fig. 2g, Fig. S2a).

Gid11 is predicted to fold into a 7-bladed β-propeller with five intrinsically disordered regions (IDR1-5) and a C-terminal tail extending from the bottom face of the β-propeller (Chen et al., 2011) (Fig. 3a, b). To understand how Gid11 interacts with the GID complex, we used AlphaFold 3 to model a GID^Gid11^ assembly consisting of full length Gid1, Gid5, Gid8, Gid11, Gid2 and Gid9, corresponding to a single substrate receptor and a single RING module. An AlphaFold 3 model of a comparable GID^Gid4^ assembly was consistent with its experimentally determined structure (Sherpa et al., 2021) (Fig. S2b-d). The GID^Gid11^ model suggested two interaction sites between Gid11 and the rest of the assembly: one between Gid1 and IDR3 of Gid11 and another involving Gid5 and the C-terminus of Gid11 (Fig. 3c, Fig. S2e, f). We tested the functional relevance of these potential interaction interfaces as follows:

– Gid4, Gid10 and Gid11 share a C-terminal Φ[DE]ΦX motif (where Φ denotes a hydrophobic amino acid) essential for their function (Kong et al., 2021; Qiao et al., 2020). Moreover, the predicted interaction between the Gid11 C-terminus and Gid5 is consistent with the experimentally observed interaction between the Gid4 C-terminus and Gid5 (Qiao et al., 2020) (Fig. S2b). In agreement with the AlphaFold 3 prediction, Gid11 variants lacking the 14 residues long C-terminal tail, lacking the last 4 residues or with a mutated Φ[DE]ΦX motif were unable to support turnover of Gpm3, Phm8 and Cpa1 in the tFT colony assay (Fig. S3a) (Kong et al., 2021). We hypothesized that the three substrate receptors should compete for binding to Gid5. Accordingly, overexpression of Gid4 or Gid10 led to stabilization of Phm8 and Cpa1 (Fig. 3d). Moreover, Hsm3, a protein with GID-dependent but Gid4/Gid10/Gid11-independent turnover (Fig. S3b), was stabilized upon overexpression of the three substrate receptors (Fig. 3d). Finally, Gid4 overexpression had no effect on Gid11 upregulation during the glucose-to-ethanol switch but blocked its subsequent degradation (Fig. 3e). This demonstrates that Gid4, Gid10 and Gid11 compete for binding to Gid5 and that the C-terminus of Gid11 is necessary for its interaction with the GID complex. Further supporting this notion, overexpression of Gid11 mutants lacking the C-terminal tail had little impact on Hsm3 turnover (Fig. S3c).
– Deletion of Gid11 IDR3 (Gid11-Δ3) but not the other predicted IDRs completely abrogated Gid11-dependent turnover of Phm8 and Cpa1 in the tFT colony assay (Fig. 3b, f). Moreover, Gid11-Δ3 overexpression had a reduced impact on Hsm3 turnover compared to overexpression of wild type Gid11 (Fig. 3g), pointing towards an impaired interaction of Gid11-Δ3 with the GID complex. The levels of the Gid11-Δ3 protein were reduced compared to wild type Gid11, but comparable to the functional Gid11-Δ1 variant (Fig. S3d). The specific sequence of IDR3 is likely required for its function as replacing IDR3 with a flexible glycine-serine linker (GGS) did not restore Gid11 expression and function (Fig. 3h, i, Fig. S3e). This suggests that Gid11 stability and interaction with the GID complex are impaired upon deletion of IDR3. However, mutating the three Gid1 determinants predicted to interact with IDR3, either alone or in combination, had no impact on turnover Phm8 and Cpa1 (Fig. 3j, Fig. S3f). We conclude that the predicted interaction between Gid11 IDR3 and Gid1 is unlikely to be functional. Taken together, these results indicate that Gid11 binds to the core GID complex via its C-terminus, competing with other Gid5-interacting substrate receptors.

**Figure 3.**
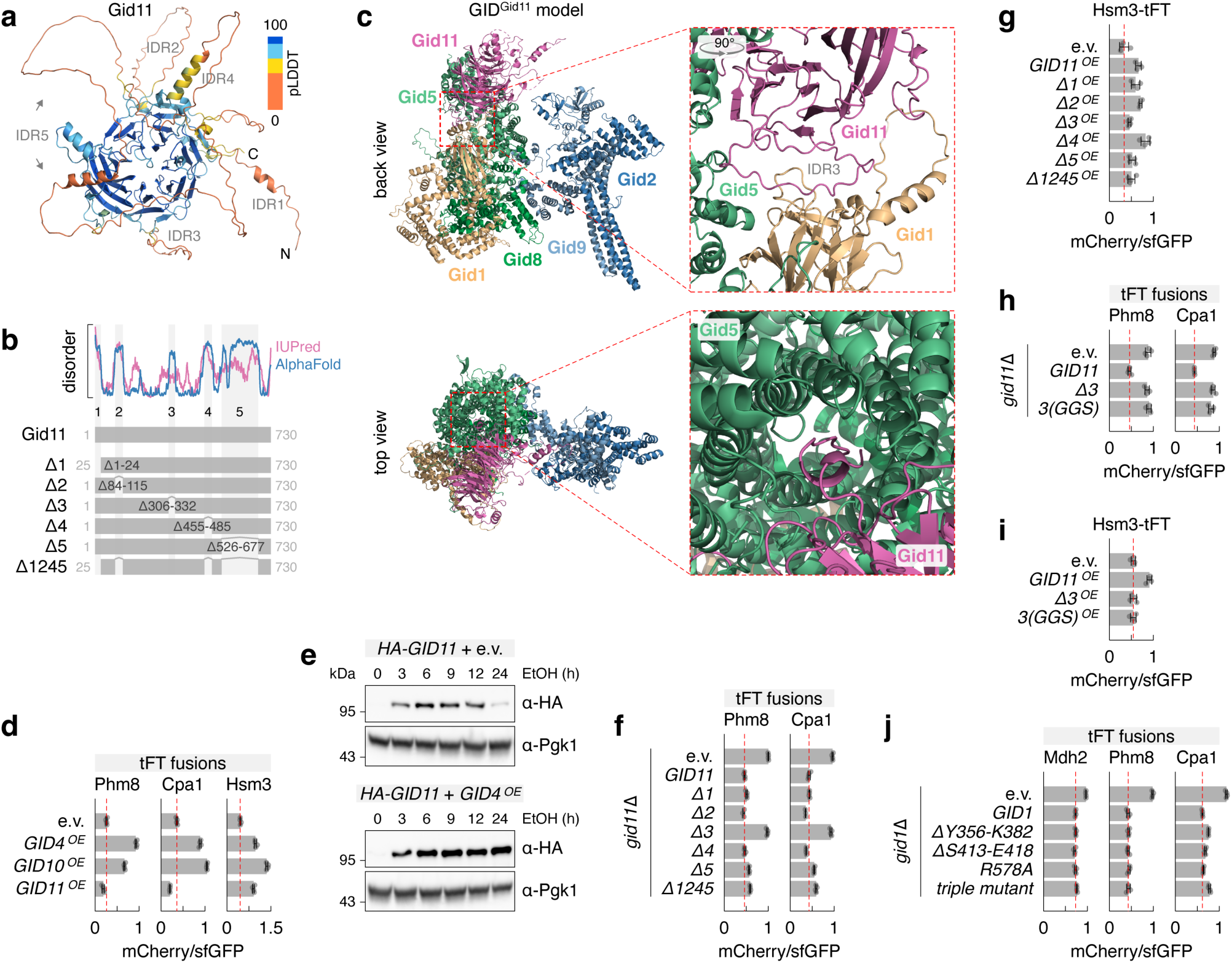
Architecture of the GID^Gid11^ complex. **a** – AlphaFold 3 model of Gid11, color coded by predicted local distance difference test (pLDDT). Five intrinsically disordered regions are marked. **b** – Cartoon of Gid11 variants lacking the five predicted intrinsically disordered regions. **c** – AlphaFold 3 model of the GID^Gid11^ assembly marked with dark outlines in Fig. 2g. Potential Gid11-Gid1 and Gid11-Gid5 interaction interfaces are highlighted (right). The following segments are omitted for clarity: Gid1 (V108-M313, H354-K382, E432-N498, R616-N676), Gid5 (P795-S868), Gid8 (M1-K25) and Gid11 (M1-D24, P83-S116, T454-S486, S525-Q678). **d** – mCherry/sfGFP ratios of colonies expressing tFT-tagged proteins (mean ± s.d., n = 4 biological replicates). e.v., empty vector; OE, overexpression. **e** – Gid11 protein levels during the glucose-to-ethanol switch. Whole-cell extracts analyzed by immunoblotting. e.v., empty vector; OE, overexpression. **f-j** – mCherry/sfGFP ratios of colonies expressing tFT-tagged proteins and HA-tagged Gid11 IDR mutants (**f-i**) (3(GGS) – IDR3 replaced with a glycine-serine linker) or 3HA-tagged Gid1 mutants (**j**) (triple mutant – ΔY356-K382 ΔS413-E418 R578A). Mean ± s.d., n = 4 biological replicates; e.v., empty vector; OE, overexpression.

To understand the functions of the GID^Gid11^ complex during the glucose-to-ethanol switch, we compared the proteomes of wild type, *gid9*Δ and *gid11*Δ strains with mass spectrometry. After the switch to ethanol, close to 30 proteins accumulated in both *gid9*Δ and *gid11*Δ backgrounds compared to the wild type (26 proteins, p-value < 0.05 and fold change (mutant/wild type) > 1.2 or 5 proteins, p-value < 0.05 and fold change (mutant/wild type) > 2) (Fig. 4a, b, Fig. S4a, b, Supplementary Data 3). This set of proteins was enriched in metabolic enzymes, 38% (10 out of 26 proteins) compared to 20% in the detected proteome (odds ratio = 2.5, p-value = 0.03 in a Fisher’s exact test) and included proteins with Gid11-dependent turnover we previously identified in screens with the tFT-tagged proteome (Gpm3, Cpa1, Yor283w, Dld3, Lys20 and Blm10, p-value < 0.05 and fold change (mutant/wild type) > 1.2) but not Phm8 or Pfk2 (Kong et al., 2021).

**Figure 4.**
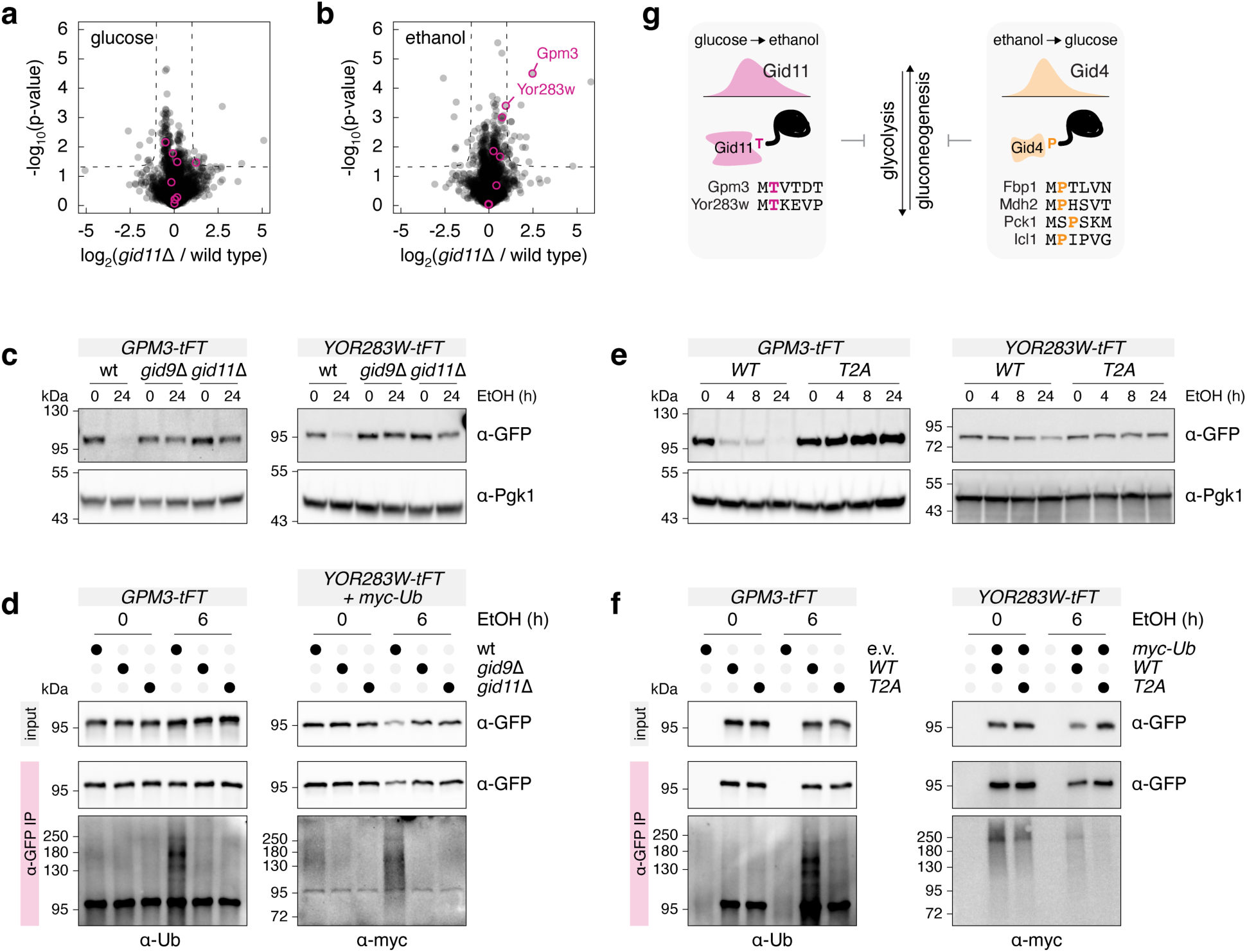
Conditional targeting of Gpm3 and Yor283w for degradation by the GID^Gid11^ complex. **a, b** – Differential proteomics of wild type and *gid11*Δ strains grown on glucose as carbon source (**a**) or 6 h after switching to ethanol (**b**). Gid11 substrates with a threonine N-terminus (Fig. 5a, with the exception of Phm8) are marked in magenta. **c** – Downregulation of Gpm3 and Yor283w during the glucose-to-ethanol switch. Whole-cell extracts analyzed by immunoblotting. **d** – Ubiquitination of Gpm3 and Yor283w during the glucose-to-ethanol switch. Whole-cell extracts (input, top) and anti-GFP immunoprecipitates (IP, bottom) analyzed by immunoblotting. Gpm3-tFT was expressed from a centromeric plasmid under the control of the *GPD* promoter, myc-tagged ubiquitin (myc-Ub) was overexpressed in *YOR283W-tFT* strains. **e** – Downregulation of Gpm3 and Yor283w, wild type (WT) and T2A mutants, during the glucose-to-ethanol switch. Whole-cell extracts analyzed by immunoblotting. **f** – Ubiquitination of Gpm3 and Yor283w, wild type (WT) and T2A mutants, during the glucose-to-ethanol switch. Whole-cell extracts (input, top) and anti-GFP immunoprecipitates (IP, bottom) analyzed by immunoblotting. Gpm3-tFT was expressed from a centromeric plasmid under the control of the *GPD* promoter, myc-tagged ubiquitin (myc-Ub) was overexpressed in *YOR283W-tFT* strains.

Although no Phm8 peptides could be detected with mass spectrometry, Phm8-tFT strongly accumulated in the absence of Gid11 during the glucose-to-ethanol switch (Fig. S4c), suggesting that Phm8 is a substrate of GID^Gid11^ during this metabolic switch. Only Pfk2-tFT, but not untagged Pfk2 or HA-tagged Pfk2, was depleted during this switch in a GID-dependent manner (Fig. S4d). This suggests that Pfk2 tagged with the large tFT moiety is a neosubstrate of GID^Gid11^. In contrast, both untagged and tFT-tagged Gpm3 and Yor283w were depleted in a GID^Gid11^-dependent manner during the glucose-to-ethanol switch (Fig. 4a-c). Ubiquitination of Gpm3 and Yor283w was substantially increased 6h after the switch to ethanol and depended on Gid9 and Gid11 (Fig. 4d, Fig. S4e). In addition, depletion and ubiquitination of both proteins were abrogated by mutation of their N-terminal threonine to alanine (Fig. 4e, f). Together, these results establish Gpm3 and Yor283w as *bona fide* substrates of the GID^Gid11^ complex during the glucose-to-ethanol switch. Gpm3 is a homolog of the Gpm1 phosphoglycerate mutase, which interconverts 3-phosphoglycerate and 2-phosphoglycerate in glycolysis and gluconeogenesis, although the functions of Gpm3 are unclear (Heinisch et al., 1998; Solis-Escalante et al., 2015). Yor283w appears to be a phosphatase with broad substrate specificity, with some similarity to Gpm1 (Ho et al., 2009; Kuznetsova et al., 2010) (Fig. S4f). We propose that while GID^Gid4^ targets for degradation gluconeogenesis enzymes during the ethanol-to-glucose switch (Chen et al., 2017; Hämmerle et al., 1998; Regelmann et al., 2003; Santt et al., 2008), GID^Gid11^ functions during the reverse glucose-to-ethanol switch, potentially modulating glycolysis by targeting for degradation Gpm3 and Yor283w (Fig. 4g).

### Specificity determinants of substrate recognition by Gid11

The observation that the N-terminal threonine in Phm8, Gpm3 and Yor283w is required for their Gid11-dependent turnover led us to propose that Gid11 targets substrates carrying threonine N-degrons (Thr/N-degrons) (Kong et al., 2021). Of note, N-terminal mutations in Blm10 and Cpa1 had no impact on their Gid11-dependent turnover (Kong et al., 2021). To define the Thr/N-degron motif recognized by Gid11, we first mutated the N-termini of all potential Gid11 substrates identified in screens with the tFT-tagged proteome (Kong et al., 2021) (Fig. 5a). Mutating the N-terminal threonine to alanine or glycine abolished Gid11-dependent turnover of seven proteins (Acs2, Dbp3, Dld3, Lys20, Pfk2, Stf2 and Top1), but not Tma10, in the tFT colony assay (Fig. 5b). This substantiates our hypothesis that Gid11 recognizes proteins with Thr/N-degrons and establishes these seven proteins (or at least their tFT fusions) as Gid11 substrates. It remains to be determined whether Blm10, Tma10 and Cpa1 are recognized by Gid11 differently or whether these three proteins are not substrates of GID^Gid11^.

**Figure 5.**
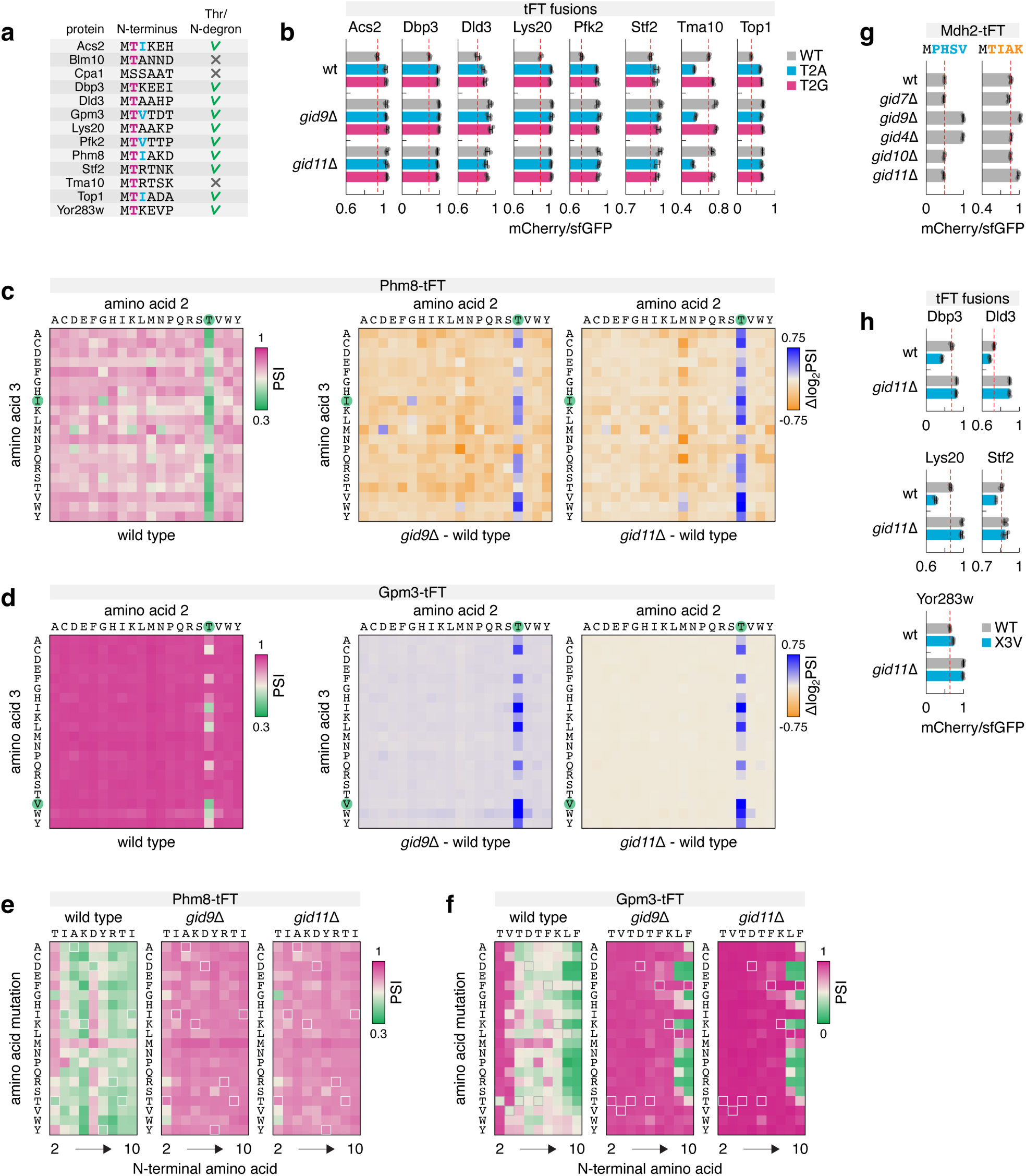
Thr/N-degron motif targeted by the Gid11 pathway. **a** – Proteins with Gid11-dependent turnover identified in a proteome-wide tFT screen (Kong et al., 2021). Gid11 substrates with a Thr/N-degron are marked (right) based on the primary sequence and functional impact of mutating the N-terminal threonine. **b** – mCherry/sfGFP ratios of colonies expressing tFT-tagged proteins, either wild type (WT) or with the second amino acid (threonine) mutated to alanine (T2A) or glycine (T2G) (mean ± s.d., n = 4 biological replicates). **c, d** – Combinatorial mutagenesis of the two N-terminal amino acids after the initiator methionine in Phm8 (**c**) and Gpm3 (**d**). Heatmaps of protein stability index (PSI, left) and differential heatmaps (right) determined by MPS profiling in wild type and *gid11*Δ backgrounds. Wild type amino acids at each position are marked in green. **e, f** – Saturation mutagenesis of Phm8 (**e**) and Gpm3 (**f**) N-termini. Heatmaps of PSI determined by MPS profiling in wild type, *gid9*Δ and *gid11*Δ backgrounds. Wild type amino acids at each position are indicated above each heatmap and highlighted with light outlines on the heatmaps. **g** – mCherry/sfGFP ratios of colonies expressing tFT-tagged wild type Mdh2 (MPHSV N-terminus) or an Mdh2 variant with the N-terminus of Phm8 (MTIAK) (mean ± s.d., n = 4 biological replicates). **h** – mCherry/sfGFP ratios of colonies expressing tFT-tagged proteins, either wild type (WT) or with the third amino acid (X) mutated to valine (X3V) (mean ± s.d., n = 4 biological replicates).

Next, we systematically mutated the N-termini of two Gid11 substrates, Phm8 and Gpm3. We constructed pooled yeast libraries expressing N-terminal variants of each tFT-tagged substrate and measured their relative turnover in wild type, *gid9*Δ and *gid11*Δ backgrounds using multiplexed protein stability (MPS) profiling (Kats et al., 2018; Kong et al., 2023a; Reinbold et al., 2023). The yeast libraries were switched from glucose to ethanol medium for 24h for these experiments. Variant libraries with all possible two amino acid combinations after the initiator methionine (positions 2 and 3) showed that threonine is the only amino acid at position 2 that allows Gid11-dependent turnover of Phm8 and Gpm3, whereas the bulky hydrophobic amino acids isoleucine, leucine, valine and tryptophan [ILVW] at position 3 preferentially promoted Gid11-dependent substrate turnover, especially in the context of Gpm3 (Fig. 5c, d, Supplementary Data 4).

Saturation mutagenesis of the nine N-terminal residues after the initiator methionine in Phm8 and Gpm3 supported this preference for a bulky hydrophobic amino acid at position 3 but revealed no further strong preferences between positions 4 and 10 (Fig. 5e, f, Supplementary Data 4). This suggests that the N-degron motif recognized by Gid11 is short, whereby the initiator methionine is followed by T[ILVW] (although the N-degron in Phm8 appears to be less restricted by the residue after threonine). Supporting this conclusion, Mdh2, a Gid4 substrate carrying a Pro/N-degron, could be converted into a Gid11 substrate by replacing its five N-terminal amino acids (MPHSV) with the N-terminus of Phm8 (MTIAK) (Fig. 5g). Moreover, 5 out of 10 Gid11 substrates have [ILVW] at position 3 (Fig. 5a), compared to 24% (124 out of 513) of MT-starting proteins in the *S. cerevisiae* proteome. The other five substrates Dbp3, Dld3, Lys20, Stf2 and Yor283w have alanine, lysine or arginine at position 3 (Fig. 5a). Mutating these amino acids to valine increased Gid11-dependent turnover in 4 out of 5 cases (Fig. 5h), further highlighting the preference for a bulky hydrophobic amino acid at position 3 in Gid11 substrates.

Methionine aminopeptidases (MetAPs) catalyze co-translational removal of the initiator methionine from nascent polypeptides typically when the second residue has a small side chain (e.g., serine, threonine, alanine, valine, cysteine, glycine, proline) (Frottin et al., 2006; Tsunasawa et al., 1985). We thus tested whether the initiator methionine is part of the Thr/N-degrons targeted by Gid11 using the ubiquitin fusion technique (Bachmair et al., 1986). In this approach, a protein of interest is expressed fused to the C-terminus of ubiquitin, followed by co-translational cleavage of the ubiquitin moiety and exposure of the next residue as the protein N-terminus (Bachmair et al., 1986) (Fig. 6a). tFT-tagged Phm8, Gpm3 and Yor283w expressed in this manner with or without the initiator methionine exhibited turnover indistinguishable from wild type proteins and were equally stabilized in *gid9*Δ and *gid11*Δ backgrounds (Fig. 6b). This indicates that the initiator methionine is not part of the Thr/N-degron.

**Figure 6.**
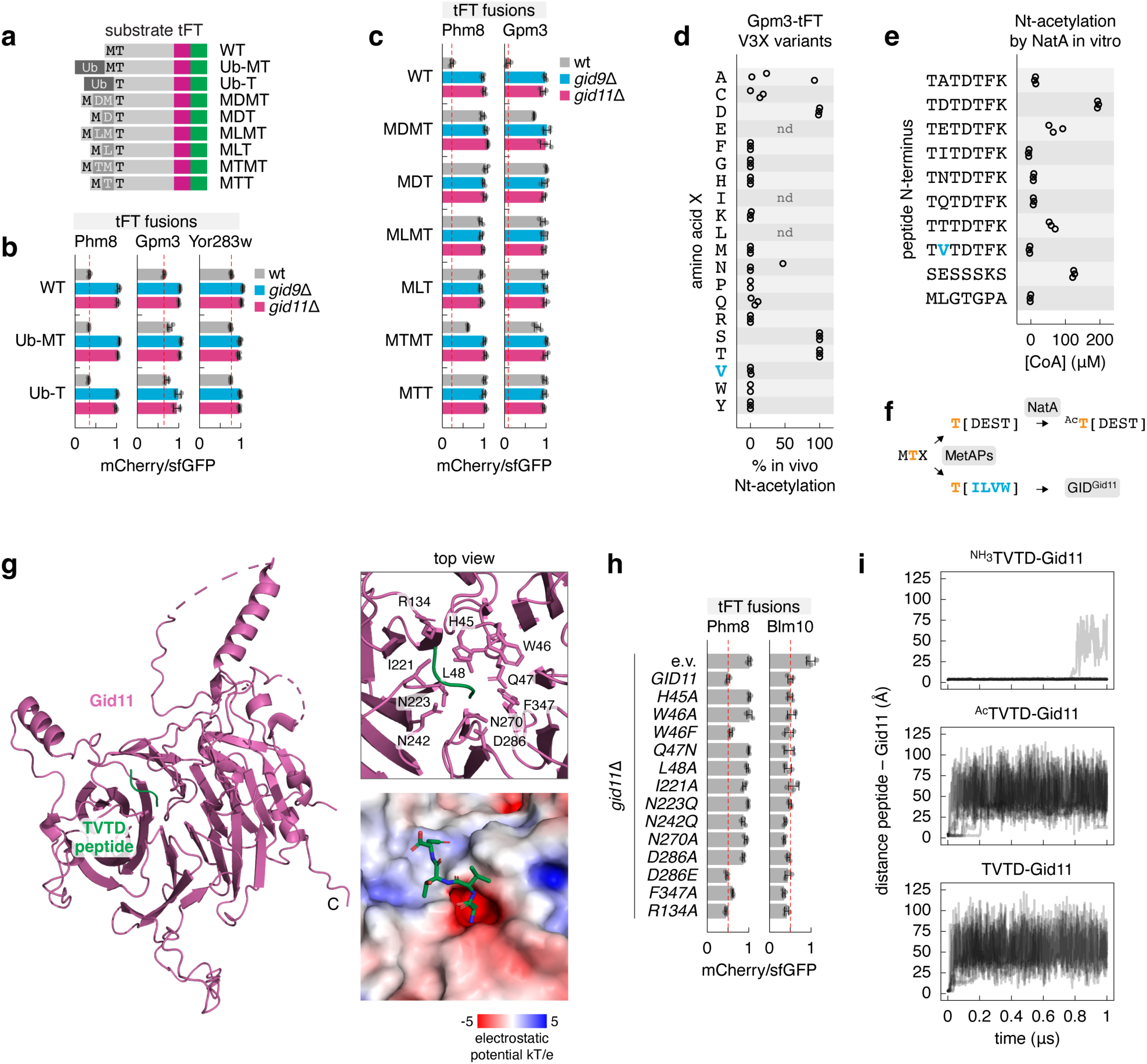
Specificity determinants of substrate recognition by Gid11. **a** – Variants of tFT-tagged Gid11 substrates with a ubiquitin (Ub) moiety fused to the N-terminus or with the indicated N-terminal mutations. **b, c** – mCherry/sfGFP ratios of colonies expressing tFT-tagged Gid11 substrates, either wild type (WT) or N-terminal variants according to **a** (mean ± s.d., n = 4 biological replicates). **d** – Nt-acetylation of Gpm3 variants with the indicated mutations of the third amino acid (valine). Gpm3-tFT variants were immunoprecipitated from strains shifted from glucose to ethanol medium for 24 h, followed by mass spectrometry analysis of N-terminal peptides (n.d., not detected; n = 3 technical replicates). **e** – N-terminal acetylation of the indicated peptides by recombinant NatA. Concentration of coenzyme A (CoA) as a readout of N-terminal acetylation (n = 3 technical replicates). **f** – Model of N-terminal processing of proteins with MTX N-termini. **g** – AlphaFold 3 model of Gid11 with the TVTD peptide corresponding to the N-terminus of Gpm3. Surface charge distribution (bottom right) and amino acids (top right) of the predicted Gid11 interaction interface. R134 is not predicted to interact with the TVTD peptide. Gid11 IDRs 1, 2, 4 and 5 are represented by dashed lines for clarity. **h** – mCherry/sfGFP ratios of colonies expressing tFT-tagged proteins and the indicated Gid11 mutants (e.v., empty vector; n = 4 biological replicates). **i** – Molecular dynamics simulations of Gid11 interaction with TVTD peptides. Simulations of tetrapeptides with the protonated N-terminus (^NH3^TVTD), acetylated N-terminus (^Ac^TVTD) or deprotonated N-terminus (TVTD). Distance between the N-terminal threonine side chain of TVTD peptides and the Thr/N-degron binding pocket of Gid11 (n = 10 replicate runs per system).

We asked whether the threonine residue needs to be exposed at the substrate N-terminus. The two yeast methionine aminopeptidases Map1 and Map2 have overlapping activities, and the double knockout is lethal (Li and Chang, 1995). Thus, instead of analyzing how the absence of MetAPs affects Gid11 substrates, we tested the impact of capping substrate N-termini by adding one or two amino acids between the initiator methionine and the threonine (Fig. 6a). Most Phm8 and Gpm3 variants with capped N-termini were fully stable (Fig. 6c), suggesting that blocking the N-terminal threonine prevents its recognition by Gid11. MTMT variants of Phm8 and Gpm3 exhibited some Gid11-dependent turnover likely because the two capping residues, threonine followed by methionine, form a weak Thr/N-degron (Fig. 6c). To understand the unexpected turnover of the MDMT variant of Gpm3, we immunoprecipitated MDMT-Gpm3-tFT from *gid9*Δ cells. Mass spectrometry analysis of the immunoprecipitates identified a peptide corresponding to the expected N-terminus, an N-terminally acetylated (Nt-acetylated) MDMTVTDTFK peptide, but also a shorter TVTDTFK peptide identical to the expected N-terminus of wild type Gpm3 (Fig. S5a). This explains the observed Gid11-dependent turnover of MDMT-Gpm3, although the origin of its N-terminally shortened version remains to be determined. Together, these results argue that the threonine residue must be exposed at the substrate N-terminus for recognition by Gid11.

After removal of the initiator methionine, threonine N-termini can be Nt-acetylated by the NatA N-terminal acetyltransferase (Aksnes et al., 2019; Arnesen et al., 2009; Mullen et al., 1989; Polevoda et al., 1999). To determine whether Gid11 substrates are Nt-acetylated, we immunoprecipitated Gpm3-tFT from *gid9*Δ cells grown on ethanol. After Nt-acetylation of unmodified N-termini with deuterated acetic anhydride *in vitro*, quantification of the relative amounts of *in vitro* and *in vivo* Nt-acetylated peptides with mass spectrometry showed that the N-terminus of Gpm3 (TVTDTFKLFILR peptide) is not Nt-acetylated *in vivo* (Fig. 6d). Moreover, compared to a NatA substrate peptide SESSSKS, a peptide with the TVTDTFK N-terminus could not be Nt-acetylated by a recombinant NatA complex *in vitro* (Fig. 6e, Fig. S5b), indicating that the Gpm3 N-terminus is not a NatA substrate. Interestingly, Gpm3 variants with the valine at position 3 mutated to acidic or hydroxylic amino acids (aspartate, glutamate, serine or threonine) were Nt-acetylated *in vivo* and the corresponding N-terminal peptides were Nt-acetylated by NatA *in vitro* (Fig. 6d, e). We conclude that Gid11 substrates are not Nt-acetylated and propose that the fate of a protein with a Thr/N-terminus is at least in part determined by the distinct preferences of NatA for acidic and hydroxylic [DEST] and Gid11 for bulky hydrophobic [ILVW] amino acids after the threonine (Fig. 6f).

To understand how Gid11 recognizes Thr/N-degrons, we used AlphaFold 3 to model Gid11 with tetrapeptides corresponding to N-termini of Gid11 substrates without the initiator methionine. In most models, the tetrapeptide was located in a negatively charged pocket on the top face of the Gid11 β-propeller, with its N-terminal threonine oriented inwards and interacting with Gid11 (Fig. 6g). We note that the model confidence (interface predicted template modeling score, ipTM) was similar for peptides starting with threonine or serine, and was largely insensitive to mutations of the second amino acid (Fig. S5c). Nevertheless, we identified ten residues in Gid11 predicted to contact the N-terminus of Gpm3 based on the model of Gid11 with the TVTD peptide (Fig. 6g). Gid11 variants with individual mutations of these amino acids, with the exception of F347, could not support Phm8 turnover in a strain lacking endogenous *GID11* (Fig. 6h). The same mutations had no impact on turnover of Blm10 (Fig. 6h), which lacks a Thr/N-degron (Fig. 5a) (Kong et al., 2021). This validates the predicted Thr/N-degron-binding pocket of Gid11.

Considering that Nt-acetylation can neutralize the positive charge of the N-terminus, we hypothesized that Nt-acetylation of Thr/N-degrons would impair their interaction with the negatively charged pocket of Gid11. Accordingly, the ipTM scores were consistently lower for Gid11 models with Nt-acetylated tetrapeptides compared to unmodified peptides (Fig. S5c, d). To further test this hypothesis, we performed molecular dynamics simulations with AlphaFold 3 models of Gid11 and an unmodified or an Nt-acetylated TVTD peptide located in the Thr/N-degron binding pocket (Fig. S5e) as a starting point. The unmodified TVTD peptide with a protonated N-terminal α-amino group and, thus, positively charged N-terminus remained in the degron-binding pocket during the 1 μs of simulation time in 9 out 10 replicates (Fig. 6i). In contrast, the Nt-acetylated peptide unbound from Gid11 in 60 ± 20 ns (mean ± s.e.m., n = 10 replicates), suggesting that Nt-acetylation reduces the affinity between Gid11 and Thr/N-degrons. Moreover, the unmodified peptide with a deprotonated and, therefore, neutral N-terminus behaved similarly to the Nt-acetylated peptide (Fig. 6i). We conclude that Nt-acetylation of Thr/N-degrons would prevent their recognition by Gid11 by neutralizing the otherwise positively charged N-terminus.

## Discussion

The GID complex was originally identified for its role in degrading gluconeogenic enzymes when yeast shift from ethanol to glucose as carbon source and transiently express the Gid4 substrate receptor (Hämmerle et al., 1998; Regelmann et al., 2003; Santt et al., 2008). Here, we establish Gid11 as a GID substrate receptor and characterize the regulation, specificity and functions of the GID^Gid11^ complex. Gid11 is induced during the reverse metabolic switch – from glucose to ethanol – where the GID^Gid11^ complex downregulates Gpm3 and Yor283w, two proteins linked to glycolysis. Unlike Gid4, which recognizes substrates via Pro/N-degrons (Chen et al., 2017; Hämmerle et al., 1998), Gid11 targets proteins that contain Thr/N-degrons. These findings illustrate how yeast cells exploit one ubiquitin ligase to control metabolic transitions in both directions by employing interchangeable substrate receptors with distinct specificities (Fig. 4g).

Similar to the Gid4 and Gid10 receptors, Gid11 binds the Gid5 subunit of the core complex with its C-terminal tail (Kong et al., 2021; Qiao et al., 2020). Indeed, the three substrate receptors can compete with one another, at least when overexpressed. This competition is reduced by conditional expression of Gid4, Gid10 and Gid11, leading to the assembly of condition- and substrate-specific GID complexes (Melnykov et al., 2019; Menssen et al., 2018, 2012; Qiao et al., 2020; Santt et al., 2008). It is possible that assembly or activity of GID complexes are further conditionally regulated by phosphorylation as multisite phosphorylation of the E2 Ubc8 was shown to anchor it to the RING module (Chrustowicz et al., 2024) and several phosphorylation sites were identified in Gid11, including in its C-terminal tail (Albuquerque et al., 2008; Lanz et al., 2021; Swaney et al., 2013).

Analysis of N-terminal processing of Gid11 substrates argues that Thr/N-degrons are constitutively exposed following co-translational removal of the initiator methionine, further emphasizing the importance of conditional Gid11 expression for timely protein degradation. Substrates bearing Thr/N-degrons bind to a shallow negatively charged pocket on Gid11. This substrate binding pocket and the C-terminal tail are on opposite faces of the Gid11 β-propeller, likely positioning the substrate for ubiquitination by the E2 engaged on the RING module (Fig. 3c). Interestingly, some proteins such as Blm10 require Gid11 for degradation but do not rely on its Thr/N-degron-binding pocket, suggesting that Gid11 may employ alternative modes of substrate recognition that remain to be explored.

Despite their threonine N-termini, Gid11 substrates are not Nt-acetylated by NatA. Whereas the Thr/N-degron motif consists of a threonine followed by a bulky hydrophobic amino acid, NatA preferentially Nt-acetylates N-terminal threonine when it is followed by acidic or hydroxylic amino acids (Fig. 6f). This preference of NatA is apparent in proteomic analyses of Nt-acetylation by yeast and human NatA (Arnesen et al., 2009). Moreover, a similar pattern of Nt-acetylation by NatA was recently observed for cysteine N-termini (Heathcote et al., 2024).

Finally, Gid11 is conserved across yeasts, including the fission yeast *Schizosaccharomyces pombe* (Kong et al., 2021). It will be interesting to determine how the specificity and functions of Gid11 changed across the ∼500 million years of evolution since fission yeasts diverged from budding yeasts (Rhind et al., 2011). Outside yeasts, other E3s, such as the UBR4 ubiquitin ligase of the Arg/N-degron pathway (Morrison et al., 2024), could target Thr/N-degrons.

## Acknowledgements

We thank Helle Ulrich (IMB, Mainz, Germany) and Ronen Marmorstein (University of Pennsylvania, Philadelphia, Pennsylvania, USA) for reagents. We thank the IMB Flow Cytometry Core Facility for the use of their instruments and cell sorting assistance; the IMB Genomics Core Facility for sequencing of MPS profiling libraries; the IMB Bioinformatics Core Facility and Frank Rühle for processing of MPS profiling sequencing data; the IMB Media Lab and the IMB Protein Production facility for support with growth media and reagents. Funding of the Deutsche Forschungsgemeinschaft (DFG, German Research Foundation) supported the BD FACSAria III SORP (P#210144599), the BD LSRFortessa SORP (P#210253511), the Next-Seq500 (P#329045328) and the Orbitrap Astral system (P#524805621). Parts of this research were conducted using the supercomputer MOGON 2 and/or advisory services offered by Johannes Gutenberg University Mainz (hpc.uni-mainz.de), which is a member of the AHRP (Alliance for High Performance Computing in Rhineland Palatinate, www.ahrp.info) and the Gauss Alliance e.V. We gratefully acknowledge the computing time granted on the supercomputer MOGON 2 at Johannes Gutenberg University Mainz (hpc.uni-mainz.de). D.A.B. was supported by the DFG in the framework of the Sonderforschungsbereich (SFB, Collaborative Research Center) 1552 (P#465145163-B04) and by the Max Planck Graduate Center with the Johannes Gutenberg University of Mainz (MPGC). V.A.C.S. was supported by the Joachim Herz Stiftung through an Add-On fellowship for Interdisciplinary Life Science. J.K.V. was supported by a Marie Sklodowska-Curie European Training Network Grant #860517 (UBIMOTIF). Research in the O.S.F. lab was partially supported by the Israel Science Foundation (ISF) founded by the Israel Academy of Sciences and Humanities (grants no. 301/2021 and 3091/23). L.S.S gratefully acknowledges support from M3ODEL and ReALity Landesforschungsinitative Rheinland-Pfalz. This work was funded by the DFG (P#557542771 to A.K., P#465145163-B04 to L.S.S., P#449991970 to K.L.) and the Research Council of Norway (FRIPRO Grant 324195 to T.A.).

## Author Contributions

Conceptualization: A.K.; experimental investigation: all authors; writing-original draft: A.K.; writing-review and editing: all authors; supervision: O.S.F., F.B., L.S.S., T.A., A.K.; funding acquisition: V.A.C.S., O.S.F., K.L., F.B., L.S.S., T.A., A.K.

## Methods

### Yeast strains and growth conditions

All yeast strains used in this work are listed in Supplementary Table 1 and are derivatives of BY4741 or Y8205. All plasmids used in this work are listed in Supplementary Table 2. Yeast genome manipulations (gene tagging and gene deletion) were performed using PCR targeting and lithium acetate transformation (Gietz and Woods, 2002; Janke et al., 2004; Knop et al., 1999). T2A mutations (Fig. 4e, f) and N-terminal capping mutations (Fig. 6c) were introduced into the *GPM3*, *PHM8* and *YOR283W* loci with the delitto perfetto approach (Storici and Resnick, 2006). Unless stated otherwise, yeast strains were grown at 30℃ in synthetic complete (SC) medium with 2% (w/v) glucose as carbon source (SC glucose) or SC glucose medium lacking histidine (SC-His glucose) for selection of plasmids based on the pRS313 and pRS413 backbones. To shift cells to other growth media, cultures grown in SC glucose or SC-His glucose media were pelleted by centrifugation, cell pellets were washed with an equal volume of water and resuspended in the new growth medium: SC ethanol (SC medium with 2% (v/v) ethanol instead of glucose), SC glycerol (SC medium with 2% (v/v) glycerol instead of glucose), carbon starvation (SC medium without a carbon source), 0.5 M NaCl (SC glucose medium with 0.5 M NaCl), nitrogen base starvation (SC glucose medium lacking nitrogen base), 1 M KH_2_PO_4_ (SC glucose medium with 1 M KH_2_PO_4_).

### tFT colony assay

Fluorescence measurements of ordered colony arrays were performed as described (Kong et al., 2023b). Briefly, yeast strains were assembled in 1536-colony format on agar plates using a pinning robot (Rotor, Singer Instruments), with 4 biological replicates per tFT strain. For each biological replicate, 4 technical replicates of the sample strain, 8 technical replicates of a control strain without a tFT and 4 technical replicates of a reference strain expressing a stable tFT construct, were arranged next to each other in 2×2 groups. Such ordered colony arrays were typically grown for 24 h at 30°C before measuring colony fluorescence.

Fluorescence measurements were performed with a multimode microplate reader equipped with monochromators for precise selection of excitation and emission wavelengths (Spark, Tecan) and a custom temperature-controlled incubation chamber. Fluorescence intensities were measured as follows: mCherry with 586/10 nm excitation, 612/10 nm emission, optimal detector gain and 40 μs integration time; sfGFP with 488/10 nm excitation, 510/10 nm emission, optimal detector gain and 40 μs integration time. For each sample strain, fluorescence intensities were first corrected for background fluorescence by subtraction of the mean of neighboring non-fluorescent colonies, and then normalized by the mean of the neighboring reference colonies to correct for spatial effects. mCherry/sfGFP ratios were subsequently calculated for each technical replicate and summarized by the mean and standard deviation per biological replicate.

### Immunoblotting

Approximately 5×10^7^ cells were harvested by centrifugation. Whole cell extracts were prepared by alkaline lysis followed by trichloroacetic acid precipitation (Knop et al., 1999; Kong et al., 2019). The precipitates were washed once with 1 ml of ethanol and resuspended in 50 μl of urea buffer (8 M urea and 10 mM β-mercaptoethanol), followed by addition of 17 μl of 4X Laemmli SDS sample buffer (250 mM Tris-HCl pH 6.8, 8% (w/v) SDS, 40% (v/v) glycerol, 5% (v/v) β-mercaptoethanol and 0.1% (w/v) bromophenol blue) and protein denaturation at 99°C for 10 min. Alternatively, the precipitates were suspended in high urea buffer (8 M urea, 200 mM Tris-HCl pH 6.8, 0.1 mM EDTA, 0.1% (w/v) bromophenol blue, 1.5% (w/v) DTT), followed by protein denaturation at 65°C for 15 min.

Whole cell extracts were then separated by SDS-PAGE, followed by immunoblotting with mouse primary antibodies against the HA epitope (dilution 1:2000, clone 12CA5, produced in-house), the myc epitope (dilution 1:2000, clone 9E10, produced in-house), GFP (dilution 1:2000, 11814460001, Roche), ubiquitin (dilution 1:1000, sc-8017, Santa Cruz Biotechnology) or Pgk1 (dilution 1:1000, 459250, Thermo Fisher Scientific) and goat anti-mouse HRP-conjugated secondary antibody (dilution 1:5000, G-21040, Thermo Fisher Scientific). Membranes were developed after addition of a chemiluminescent horseradish peroxidase (HRP) substrate (SuperSignal West Pico PLUS, 34580, Thermo Fisher Scientific) and imaged using a ChemiDoc MP system (Bio-Rad).

### Luciferase assay

Nano-luciferase activity was measured using the Nano-Glo Live Cell Assay System (N2011, Promega). All measurements were performed in white opaque 96-well microtiter plates (OptiPlate-96, 6005290, PerkinElmer). Each well contained 100 μl of yeast culture and 25 μl of the reaction mix with the substrate furimazine. Signal intensities were quantified immediately after furimazine addition with a plate reader (Infinite 200 PRO, Tecan). After subtraction of the background signal determined from a strain without luciferase expression, signal intensities were normalized by cell concentration (optical density at 600 nm, OD_600_) and summarized by the mean and standard deviation of biological replicates.

### Quantification of transcript levels with RT-qPCR

Total RNA was extracted from approximately 5×10^7^ cells using the hot phenol method as described (Kong et al., 2019). Contaminating genomic DNA was removed using the RNase-Free DNase Set (79254, Qiagen), followed by reverse transcription with the SuperScript III First-Strand Synthesis System (18080051, Thermo Fisher Scientific) according to manufacturer’s instructions. Quantitative PCR was performed using the PowerUp SYBR Green Master Mix (A25780, Thermo Fisher Scientific) and primers Gid11_F/R (primers #1 and #2 in Supplementary Table 3) for detecting the *GID11* mRNA or primers Alg9_F/R (primers #3 and #4 in Supplementary Table 3) for detecting the *ALG9* mRNA as a stable reference (Teste et al., 2009). All samples were analyzed in triplicates. The 2^-ΔΔCt^ method was used to calculate the levels of *GID11* mRNA relative to *ALG9*.

### Genome-wide screens for regulators of GID11 expression

Query strains with chromosomally integrated constructs expressing mCherry and sfGFP under control of the *GID11* or *TDH3* promoters were crossed with an arrayed library of haploid strains carrying deletion alleles of most non-essential genes (Giaever et al., 2002; Winzeler et al., 1999). Mating, diploid selection, sporulation and haploid selection were performed in 1536-colony format following the SGA (synthetic genetic array) methodology (Baryshnikova et al., 2010; Tong et al., 2001) by sequential pinning of yeast colonies on agar plates with appropriate selective media using a pinning robot (Rotor, Singer Instruments). Each cross was performed in 4 technical replicates arranged next to each other in a 2×2 group. The resulting haploid colony arrays were grown for 24 h on SC agar plates lacking leucine.

mCherry and sfGFP fluorescence intensities of colonies were measured with a multimode microplate reader (Spark, Tecan) as described above, but with two fixed detector gains to extend the detection range. The colony arrays were also photographed to identify failed crosses based on colony size. After excluding measurements from failed crosses, fluorescence intensities were log-transformed, corrected for spatial effects and normalized to the median per screen plate (Fung et al., 2022). Technical replicates were summarized by the mean and standard deviation. A two-sided t-test was used to compare each sample to the screen median and to compute p-values, adjusted for multiple testing using the method of Benjamini-Hochberg.

### Immunoprecipitation of tFT-tagged proteins

Approximately 10^9^ cells were harvested by centrifugation, resuspended in IP buffer (50 mM Tris-HCl pH 7.5, 100 mM NaCl, 1% (v/v) Triton X-100, 2 mM EDTA and protease inhibitors (4693159001, Sigma-Aldrich)) and lysed by vortexing at 4°C with acid-washed glass beads. Lysates were clarified by centrifugation at 16000 to 21000 g. Clarified protein lysates were incubated with Dynabeads M-280 sheep anti-rabbit IgG (11203D, Thermo Fisher Scientific) pre-coated with an anti-GFP antibody (ab290, Abcam). After incubation for 2 h at 4°C with gentle rocking, the beads were washed three times with IP buffer, and the bound proteins were eluted and denatured with IP buffer supplemented with the appropriate amount of 4X Laemmli SDS sample buffer (250mM Tris-HCl pH 6.8, 8% (w/v) SDS, 40% (v/v) glycerol, 5% (v/v) β-mercaptoethanol and 0.1% (w/v) bromophenol blue) at 99°C for 10 min. Protein samples were then analyzed by immunoblotting.

Alternatively, cells were resuspended in IP buffer (25 mM Tris-HCl pH 8, 150 mM NaCl, 0.5% (w/v) sodium deoxycholate, 1% (v/v) Triton X-100, 0.1% (w/v) SDS, 1 mM EDTA and protease inhibitors (4693159001, Sigma-Aldrich)). Clarified protein lysates were incubated with magnetic agarose beads coated with GFP nanobodies (Fridy et al., 2014) (produced in-house). The bound proteins were eluted and denatured with elution buffer (25 mM Tris-HCl pH 8, 1% (w/v) SDS) at 90°C for 10 min with 1500 rpm shaking. Protein samples were then processed for mass spectrometry.

### Gid1-HA and HA-Gid11 immunoprecipitation followed by mass spectrometry

Clarified whole cell lysates were prepared from approximately 10^9^ cells as described above for tFT-tagged strains and incubated with Dynabeads pre-coated with an anti-HA antibody (ab9110, Abcam). The bound proteins were eluted and denatured with IP buffer supplemented with the appropriate amount of 4X NuPAGE LDS Sample Buffer (NP0007, Thermo Fisher Scientific) at 99°C for 10 min and processed as detailed below.

### Mass spectrometry analysis of whole cell proteomes

Approximately 2×10^8^ cells were lysed in MS buffer (25mM Tris-HCl pH 8.0, 150mM NaCl, 1% (v/v) Triton X-100, 0.5% (w/v) sodium deoxycholate, 0.1% (w/v) SDS, 1mM EDTA and protease inhibitors (4693159001, Sigma-Aldrich)) by vortexing at 4°C in the presence of acid-washed glass beads. The lysates, clarified by centrifugation, were diluted to 1 μg/μl protein concentration, and 50 μl of each sample were processed for mass spectrometry.

Samples were separated on a 4-12% NOVEX NuPAGE gradient SDS gel (Thermo Fisher Scientific) for 10 min at 180 V in 1X MES buffer (Thermo Fisher Scientific). Proteins were fixed and stained with Coomassie G250 brilliant blue (Carl Roth). The gel lanes were cut, and each lane was minced into 1×1 mm pieces. Gel pieces were destained with a 50% ethanol/50 mM ammonium bicarbonate (ABC) solution. Proteins were reduced in 10 mM DTT (Sigma-Aldrich) for 1 h at 56°C and then alkylated with 50 mM iodoacetamide (Sigma-Aldrich) for 45 min at room temperature. Proteins were digested with mass spectrometry grade trypsin (Serva) overnight at 37°C. Peptides were extracted from the gel by two incubations with 30% ABC/acetonitrile and three subsequent incubations with pure acetonitrile. The acetonitrile was finally evaporated in a concentrator (Eppendorf) and samples were loaded on StageTips (Rappsilber et al., 2007) for desalting and storage. For mass spectrometric analysis on the Q Exactive platform, peptides were separated on a 30 cm self-packed column with a 75 μm inner diameter filled with ReproSil-Pur 120 C18-AQ (Dr. Maisch GmbH) mounted to an EASY HPLC 1000 (Thermo Fisher Scientific) and sprayed online into a Q Exactive Plus mass spectrometer (Thermo Fisher Scientific). We used a 94 min gradient from 2% to 40% acetonitrile in 0.1% formic acid at a flow of 225 nl/min. The mass spectrometer was operated with a top 10 MS/MS data-dependent acquisition scheme per MS full scan.

For mass spectrometric analysis on the Exploris platform, peptides were separated on a 50 cm self-packed column with a 75 μm inner diameter filled with ReproSil-Pur 120 C18-AQ (Dr. Maisch GmbH) mounted to an EASY HPLC 1200 (Thermo Fisher Scientific) and sprayed online into an Exploris 480 mass spectrometer (Thermo Fisher Scientific). We used a gradient from 2% to 40% acetonitrile in 0.1% formic acid at a flow of 250 nl/min with a duration of 75 min (immunoprecipitation samples) and 105 min (whole cell proteomes). The mass spectrometer was operated with a top 15 (immunoprecipitation samples) and top 20 (whole cell proteomes) MS/MS data-dependent acquisition scheme per MS full scan.

Mass spectrometry raw data were searched using the Andromeda search (Cox et al., 2011) integrated into MaxQuant suite 1.6.5.0 and 1.6.10.43 (Cox and Mann, 2008) using the UniProt *Saccharomyces cerevisiae* database (6649 entries). In all analyses, carbamidomethylation at cysteine was set as fixed modification while methionine oxidation and protein N-acetylation were considered as variable modifications. Match between run option was activated. Reverse hits, proteins only identified by site, protein groups based on one unique peptide, and known contaminants were removed. The LFQ (label-free quantitation) values were log_2_-transformed and the median across replicates was calculated. This enrichment was plotted against −log_10_-transformed p-values (Welch t-test).

### Mass spectrometry analysis of *in vivo* protein N-terminal acetylation

#### *In vitro* protein acetylation and enzymatic digestion

tFT-tagged immunoprecipitates eluted from the beads were reduced with DTT, followed by alkylation by iodoacetamide, quenching by DTT, and purification using the SP3 approach (Hughes et al., 2019). Thereafter, the purified proteins were eluted twice in 50 μl of 3 M guanidinium chloride, 250 mM MOPS pH 7.9, at 37°C with orbital shaking for 10 min. *In vitro* protein acetylation reaction was then initiated by adding D6-acetic anhydride (175641, Sigma-Aldrich) to 50 mM followed by incubation at 37°C with orbital shaking for 30 min. The reaction was repeated once. Afterwards, unreacted D6-acetic anhydride was quenched by adding ammonium bicarbonate buffer. This step also diluted the concentration of guanidinium chloride to 1 M. Following incubation at 37°C with orbital shaking for 10 min, proteins were digested by trypsin (2 μg per sample) at 37°C overnight. The resultant peptide solution was acidified with formic acid and purified by solid phase extraction in C18 StageTips (AttractSPE Bio - C18, Affinisep) (Rappsilber et al., 2003).

#### Liquid chromatography tandem mass spectrometry

Peptides were separated via an in-house packed 45 cm analytical column (inner diameter 75 μm; ReproSil-Pur 120 C18-AQ 1.9-μm silica particles, Dr. Maisch GmbH) on a Vanquish Neo UHPLC system (Thermo Fisher Scientific). The online reversed-phase chromatography separation was conducted through a 70 min non-linear gradient of 1.6-32% acetonitrile in 0.1% formic acid at a nanoflow rate of 300 nl/min. The eluted peptides were sprayed directly by electrospray ionization into an Orbitrap Astral mass spectrometer (Thermo Fisher Scientific). Mass spectrometry was conducted in data-dependent acquisition mode using a top50 method with one full scan in the Orbitrap analyzer (scan range 325 to 1300 m/z, resolution 120000, target value 3×10^6^, maximum injection time 25 ms) followed by 50 fragment scans in the Astral analyzer via higher energy collision dissociation (HCD; normalized collision energy 26%, scan range 150 to 2000 m/z, target value 1×10^4^, maximum injection time 10 ms, isolation window 1.4 m/z). Precursor ions of unassigned, +1 or higher than +6 charge state were rejected. Additionally, precursor ions already isolated for fragmentation were dynamically excluded for 15 s.

#### Mass spectrometry data processing

Mass spectrometry raw data files were processed using MaxQuant software (version 2.1.3.0) (Cox and Mann, 2008). MS/MS mass spectra were searched using Andromeda search engine (Cox et al., 2011) against a target-decoy database containing the forward and reverse protein sequences of UniProt *Saccharomyces cerevisiae* reference proteome (6089 entries, release 2022_03), the N-terminal variants of the Gpm3 protein and a default list of common contaminants. Trypsin/P specificity was chosen. A maximum of 2 missed cleavages was tolerated. Cysteine carbamidomethylation was set as fixed modification. Methionine oxidation, protein N-terminal acetylation, D3-acetylation at lysine, serine, threonine and tyrosine residues as well as the protein N-terminus were assigned as variable modifications. Up to 6 modifications per peptide were allowed. The minimum peptide length was set to 7 amino acids. The “second peptides” option was switched on. The “match between runs” function was turned off. False discovery rate (FDR) was set to 1% at both peptide and protein levels. To compare the levels with or without *in vivo* acetylation at the N-terminus of the reporter protein, all detected N-terminal peptide sequences of the reporter protein were extracted from the MaxQuant output modificationSpecificPeptides.txt file. Among these peptide sequences, the ones that contained an N-terminal wild type acetylation modification were considered as *in vivo* Nt-acetylation. The remaining ones that were detected with N-terminal D3-acetylation or no acetylation at the N-terminus were considered as without *in vivo* Nt-acetylation. The intensities of these peptides were then summed separately for the ones with or without *in vivo* Nt-acetylation.

### Structural modeling with AlphaFold

Protein sequences were retrieved from the Saccharomyces Genome Database (Engel et al., 2025) (SGD, https://yeastgenome.org). Structural models of Gid11, GID^Gid4^, GID^Gid11^ and Gid11-tetrapeptide complexes were generated with AlphaFold 3 (Abramson et al., 2024), using the AlphaFold server (https://alphafoldserver.com/) or an internal Nextflow-based pipeline (https://github.com/imbforge/fold2go) with AlphaFold3 model weights released by (Abramson et al., 2024).

Input sequences were formatted as JSON files, with Nt-acetylated threonine and serine indicated with the CCD (chemical component dictionary) codes THC and SAC, respectively. For each model, 20 independent structural predictions were generated using 20 different seeds to capture conformational variability. The top prediction was selected based on the highest interface predicted template modeling (ipTM) and predicted template modeling (pTM) scores, provided in the AlphaFold 3 output. When multiple predictions shared identical top scores, the first occurrence was selected for downstream analysis.

### Multiplexed protein stability profiling

Analysis of Phm8-tFT and Gpm3-tFT N-terminal variant libraries by MPS profiling involved construction of libraries, fluorescence-activated cell sorting based on the tFT readout, DNA extraction from the sorted populations and preparation of amplicon libraries, followed by deep sequencing of the amplicon libraries and analysis of the sequencing data as described (Kong et al., 2023b).

Briefly, pooled libraries were constructed by homologous recombination in yeast using degenerate oligonucleotides or oligonucleotide pools (Kats et al., 2018; Kong et al., 2023a; Reinbold et al., 2023). All libraries, except for the Phm8 saturation mutagenesis library, were assembled using degenerate oligonucleotides. These oligonucleotides (#5-8 and #12-29 in Supplementary Table 3) were designed with the variable region flanked with 37 nucleotide overhangs homologous to the insertion site in the pKEK184 and pKEK447 vectors. Complementary degenerate oligonucleotides were annealed at 20 μM total concentration in a total volume of 40 μl. The Phm8 saturation mutagenesis library was synthesized as an oligonucleotide pool (#9) with 18 and 23 nucleotide overhangs. The pool was amplified by PCR to extend the overhangs to 37 nucleotides: 0.8 pmol of the pool as template, with primers Phm8_extend_F/R (#10 and #11 in Supplementary Table 3), 10 cycles. Annealed degenerate oligonucleotides or PCR-amplified pools (entire reaction volume) were subsequently combined with 15 μg of SalI-linearized pKEK184 or pKEK447 vectors and transformed into approximately 2.4×10⁹ competent yeast cells with the desired genotype. For both types of libraries, the transformed cells were cultured for 2 days until saturation (OD600 > 7) and stored as glycerol stocks at −80°C.

For sorting, libraries were inoculated from frozen stocks into SC-His glucose medium and grown overnight to saturation, diluted into fresh medium to 0.2 OD_600_, grown to 0.8 OD_600_. and then shifted to SC-His ethanol medium for 24 h. Fluorescence activated cell sorting was performed on a FACSAria III SORP cell sorter (BD Biosciences) with a 70 μm nozzle (70 psi) using a 561 nm laser for mCherry excitation, a 600 nm long pass mirror and a 610/20 nm band pass filter for mCherry detection, a 488 nm laser for sfGFP excitation, a 505 nm long pass mirror and a 530/30nm band pass filter for sfGFP detection. For each library, 5×10^6^ cells were sorted according to the mCherry/sfGFP ratio into 8 stability bins of approximately equal width on the mCherry/sfGFP scale (Fig. S6a-d).

The sorted populations were expanded, followed by DNA extraction from 2×10^8^ cells (QIAprep spin miniprep kit, 27106, Qiagen). The extracted DNA was used as a template in a two-step PCR with a high-fidelity DNA polymerase (NEBNext ultra II Q5 master mix, M0544L, NEB) to amplify the variable region of interest, introduce bin and sample barcodes and the P5/P7 adaptors (primers #30-99 for PCR1, primers #100-101 for PCR2). The resulting amplicon libraries were sequenced on a NextSeq 500 system (Illumina) with a 150-cycle kit in 2×75bp paired-end mode.

Analysis of sequencing data, followed by calculation of protein stability indices (PSIs) per sequence variant were calculated from read count data as described (Kong et al., 2023b) (https://github.com/Khmelinskii-Lab/Das1_C-degrons/tree/main/NGSpipe2go-MPSprofiling). Briefly, PSIs were determined on read counts per amino acid sequence generated with the Biostrings R package. First, the observed read counts per bin for each sequence variant were normalized by the summed read counts for all sequence variants of this bin and by the corresponding cell fraction count. The PSI was then calculated as the sum of these values over all bins multiplied with the corresponding cell fraction indices divided by the sum over all bins. Downstream analysis and data visualization were performed in R (R Core Team, 2020). Technical replicates were combined using linear regression (lmFit) in the limma R package (Ritchie et al., 2015).

### *In vitro* N-terminal acetylation assay

A NatA complex consisting of yeast Naa10/Ard1 residues 1-226 and full-length yeast Naa15/Nat1 with an N-terminal 6xHis tag was expressed and purified essentially as described (Deng et al., 2019). Briefly, plasmids encoding the two proteins were co-transformed into *Escherichia coli* BL21 cells. For protein expression, the transformed bacteria were grown in shake flask culture with Terrific Broth at 37°C until an optical density OD_600_ of 0.5 was reached, cooled rapidly, induced with 0.1 mM IPTG, and cultured overnight at 18°C. The next day, cells were harvested by centrifugation and resuspended in buffer containing 50 mM Tris-HCl pH 8, 0.3 M NaCl, 10 mM imidazole, 0.5 mM tris(2-carboxyethyl)phosphine (TCEP) and protease inhibitor cocktail. The cell suspension was flash frozen in liquid nitrogen and stored at −80°C.

For protein purification, the cells were thawed, lysed by sonication, and the lysate was clarified by centrifugation. The clarified lysate was incubated with nickel-nitrilotriacetic acid matrix, the matrix was washed with buffer containing 50 mM Tris-HCl pH 8, 0.5 M NaCl, 20 mM imidazole, and 0.1 mM TCEP, and bound proteins were eluted with the same buffer but containing 0.3 M imidazole. Elution fractions with the NatA complex were dialyzed overnight against buffer containing 25 mM Na-citrate pH 5.5 and 0.1 mM TCEP. The dialyzed sample was applied to Hitrap SP HP column (Cytiva), the column was washed with same buffer containing 0.1 M NaCl, and bound proteins were eluted with a gradient from 0.1 to 1 M NaCl. The peak containing yeast NatA was pooled, concentrated and applied to a Superdex 200 column (Cytiva) equilibrated with buffer containing 25 mM HEPES pH 7, 0.2 M NaCl, and 1 mM TCEP. The elution fractions containing the NatA complex were pooled, concentrated to approximately 1.5 mg/ml, aliquoted, flash frozen in liquid nitrogen, and stored at −80°C until use.

Acetylation activity of the purified NatA complex against peptides was measured as described (Foyn et al., 2017). The peptides consisted of 7 specific amino acids followed by a common 17 amino acids long stretch (RWGRPVGRRRRPVRVYP) to ensure solubility: SESSSKS (positive control, NatA substrate), MLGTGPA (negative control) and a series of peptides based on the Gpm3 N-terminus (TVTDTFK and variants with the valine replaced with alanine, aspartate, glutamate, isoleucine, asparagine, glutamine or threonine) (Innovagen AB). The reaction buffer consisted of 50 mM HEPES pH 7.5, 100 mM NaCl and 1 mM EDTA. 50 μl reactions containing 0.2 mM peptide, 0.2 mM acetyl-coenzyme A and 0.2 μM of recombinant NatA complex in reaction buffer were incubated for 30 min at 37°C. After 30 min, the reactions were quenched by adding 100 μl of 100 mM NaH_2_PO_4_ pH 6.8 and 3.2 M guanidine-HCl. Identically incubated reactions where the enzyme was added only after quenching were used as controls. 5,5′-dithiobis-(2-nitrobenzoic acid) (DTNB) was added, and absorbance at 412 nm was measured with a Tecan Infinite M Nano plate reader (Tecan). Coenzyme A concentrations were calculated from the absorbance after pathlength correction using the extinction coefficient 13700 M^-1^ cm^-1^.

### Molecular dynamics simulations

The starting structures for the molecular dynamics (MD) simulations were two AlphaFold 3 models of Gid11-Δ1245 (Gid11 hereafter) complexed with the unmodified and the Nt-acetylated TVTD peptide located in the Thr/N-degron binding pocket (Fig. S5e). As is common practice when simulating Nt-acetylated protein residues (Lemkul, 2024), the Nt-acetylated threonine residue was modelled as two separate residues: an acetyl group, with CCD code ACE, followed by a standard threonine residue, with CCD code THR. To accomplish this, the starting structure was manipulated with PDBFixer 1.11 (Eastman et al., 2017) (https://github.com/openmm/pdbfixer/releases/tag/v1.11). To conclude the preparation of the input structures, hydrogen atoms were added to the heavy-atom-only models with PDB2PQR 3.7.1 (Dolinsky et al., 2004) (https://github.com/Electrostatics/pdb2pqr/releases/tag/v3.7.1), which integrates the software for protein pK_a_ estimation PROPKA 3 (3.5.1) (https://github.com/jensengroup/propka/releases/tag/v3.5.1) (Olsson et al., 2011; Søndergaard et al., 2011). With this software, appropriate protonation states were assigned to the titratable groups in the complexes of Gid11 with the unmodified and Nt-acetylated TVTD (^Ac^TVTD hereafter) peptides assuming pH 6.5. For the N-terminus of the unmodified TVTD peptide, both its protonated (^NH3^TVTD hereafter) and deprotonated (TVTD hereafter) forms were considered to test the hypothesis that the neutralization of the positive charge of the terminus impairs the interaction with the negatively charged pocket of Gid11.

Starting from the generated input structures of the complexes ^NH3^TVTD-Gid11, ^Ac^TVTD-Gid11 and TVTD-Gid11, all-atom MD simulations were performed with GROMACS 2022.5 (Bauer et al., 2023; Van Der Spoel et al., 2005). The July 2022 version of the port for GROMACS of the force field CHARMM36m (https://mackerell.umaryland.edu/charmm_ff.shtml#gromacs), a version of the CHARMM force field optimized for the simulation of proteins (Best et al., 2012; Huang et al., 2017; Huang and MacKerell, 2013), was used in conjunction with the CHARMM-modified version of the water model TIP3P (Jorgensen et al., 1983) included in the port.

The following steps constitute a standard MD simulation workflow with GROMACS. Firstly, the systems were prepared for the dynamics runs: the tetrapeptide-Gid11 complexes were centered inside rhombic-dodecahedral simulation boxes (with edges at least 1.2 nm away from the molecules), the complexes were then solvated in water and Na^+^ and Cl^−^ ions were introduced at a concentration of 0.15 M to neutralize the charges of the proteins. At the end, the total number of atoms was around 10^5^ for the three systems. To conclude the preparation, energy minimization was conducted to optimize the geometry of the molecules and reduce steric clashes, especially of the newly introduced water molecules and ions. Secondly, for each tetrapeptide-Gid11 system three consecutive dynamics runs were performed: an NVT equilibration of 1 ns to attain the target temperature of 300 K, an NPT equilibration of 4 ns to attain the target pressure of 1 bar and a production run under the NPT (constant number of particles, pressure and temperature) ensemble with 10 replicates (Knapp et al., 2018) of 1 µs each. One of the 10 replicates was initiated with the velocities inherited from the NPT equilibration whereas the remaining 9 with new random velocities drawn from a Maxwell-Boltzmann distribution at 300 K. In the three dynamics runs, the leapfrog integrator with a time-step size of 2 fs (Sahil et al., 2023), the velocity-rescaling thermostat (Bussi et al., 2007) and the cell-rescaling barostat (Bernetti and Bussi, 2020) were used. Other simulation parameters included the recommendations for GROMACS simulations with CHARMM36 (Bauer et al., 2023) (full sets of parameters in Supplementary Data 5).

From the simulated molecular trajectories, the distance between the tetrapeptides and Gid11, defined as the distance between the center of geometry (CoG) of the side chain of the N-terminal threonine of the tetrapeptides and the CoG of the side chains of all the residues in the Thr/N-degron binding pocket (Fig. 6h), as a function of time was computed (Fig. 6i). The mean lifetimes of the tetrapeptides in the binding pocket were calculated by categorizing the state of the tetrapeptide-Gid11 systems at each time step as “bound” or “unbound” using the method of transition-based state assignment (Buchete and Hummer, 2008). The trajectory files were processed with the library MDAnalysis 2.8.0 (Gowers et al., 2016; Michaud-Agrawal et al., 2011) (https://github.com/MDAnalysis/mdanalysis/releases/tag/package-2.8.0), the numerical computations on the trajectories were performed with the library NumPy 2.2.1 (Harris et al., 2020) (https://github.com/numpy/numpy/releases/tag/v2.2.1), and the distance plots (Fig. 6i) were made with the library Matplotlib 3.10.0 (Hunter, 2007) (https://github.com/matplotlib/matplotlib/releases/tag/v3.10.0) in Python 3.11.11.

### Statistics and reproducibility

SGA screens were performed in four technical replicates (four colonies of the same strain) placed next to each other on the screen plates. In MPS profiling experiments, library complexity was ensured by analyzing a number of cells at least 50–100 fold larger than the number of sequence variants in a library. One to three technical replicates corresponding to independent sorting and sequencing of the same yeast library were analyzed, as all variants of each peptide could be directly compared. All other yeast experiments were performed with at least two biological replicates, i.e., independent clones isolated from a single transformation, as is standard in the field.

## Supplementary Figures

**Figure S1.**
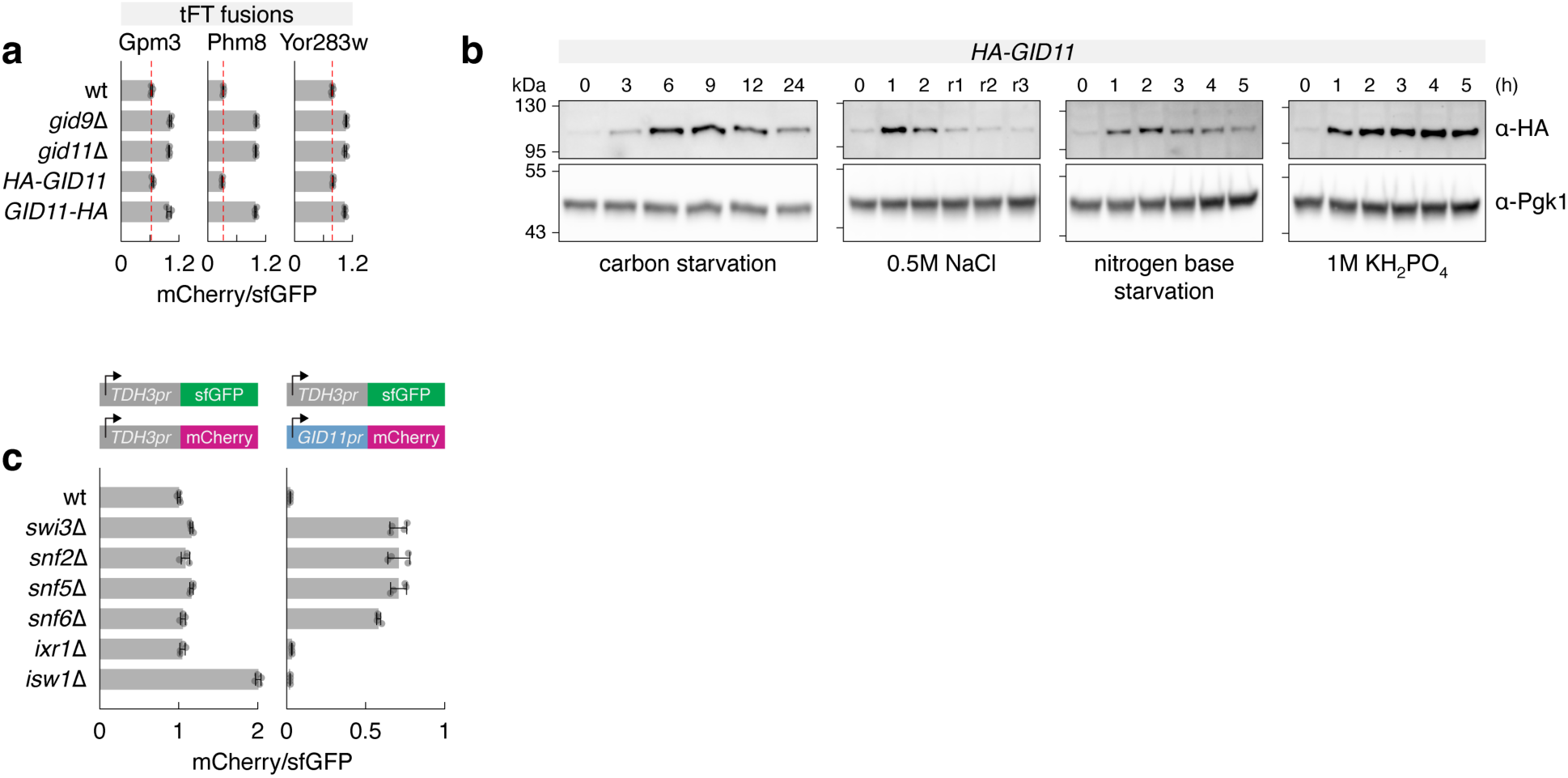
**a** – mCherry/sfGFP ratios of colonies expressing tFT-tagged proteins (mean ± s.d., n = 4 biological replicates; wt, wild type). Gid11 was tagged with the HA epitope tag at the endogenous chromosomal locus. **b** – Gid11 protein levels during the switch from glucose to medium lacking a carbon source (carbon starvation) or nitrogen base (nitrogen base starvation), or after switching to glucose medium supplemented with 0.5 M NaCl or 1 M KH_2_PO_4_. r1, r2 and r3, 1-3 h of recovery after return to glucose medium. Whole-cell extracts analyzed by immunoblotting. **c** – mCherry/sfGFP ratios of colonies expressing the indicated pairs of reporters (mean ± s.d., n = 4 biological replicates).

**Figure S2.**
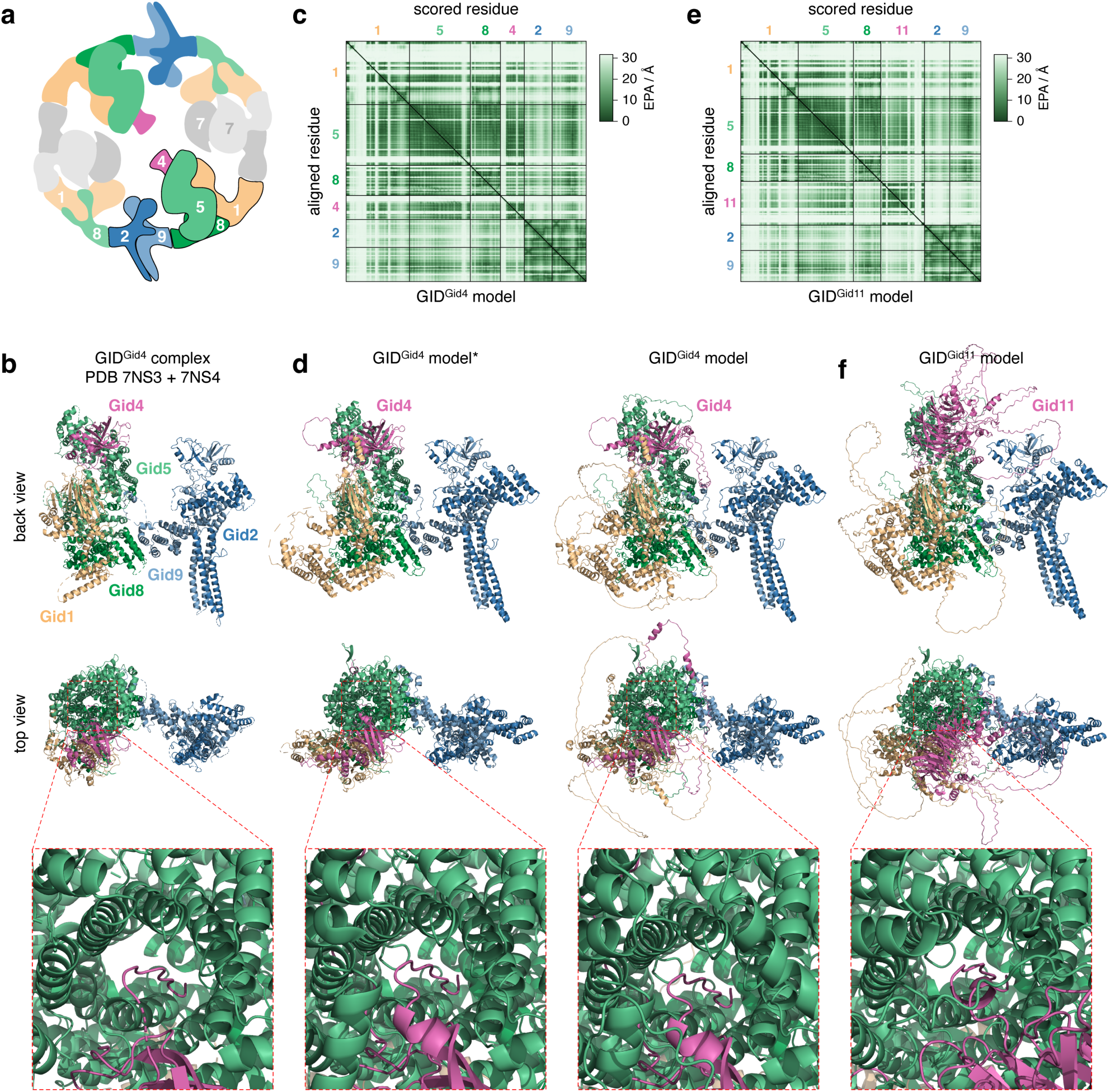
**a** – Cartoon of the GID^Gid4^ ring complex, adapted from (Sherpa et al., 2021). **b** – Structure of the GID^Gid4^ complex marked with dark outlines in **a** (PDB: 7NS3 and 7NS4). **c-f** – AlphaFold 3 models of the GID^Gid4^ assembly marked with dark outlines in **a** (**c**, **d**) and GID^Gid11^ assembly (Fig. 3c). Heatmaps of expected predicted error (EPA) (**c**, **e**). GID subunits are colored according to **a**. Predicted Gid4-Gid5 and Gid11-Gid5 interactions are highlighted (**d**, **f**, bottom). *, the following segments in the GID^Gid4^ model are omitted for clarity: Gid1 (V108-M313, H354-K382, E432-N498, R616-N676), Gid5 (P795-S868), Gid8 (M1-K25), Gid4 (M1-S89) (**d**, left).

**Figure S3.**
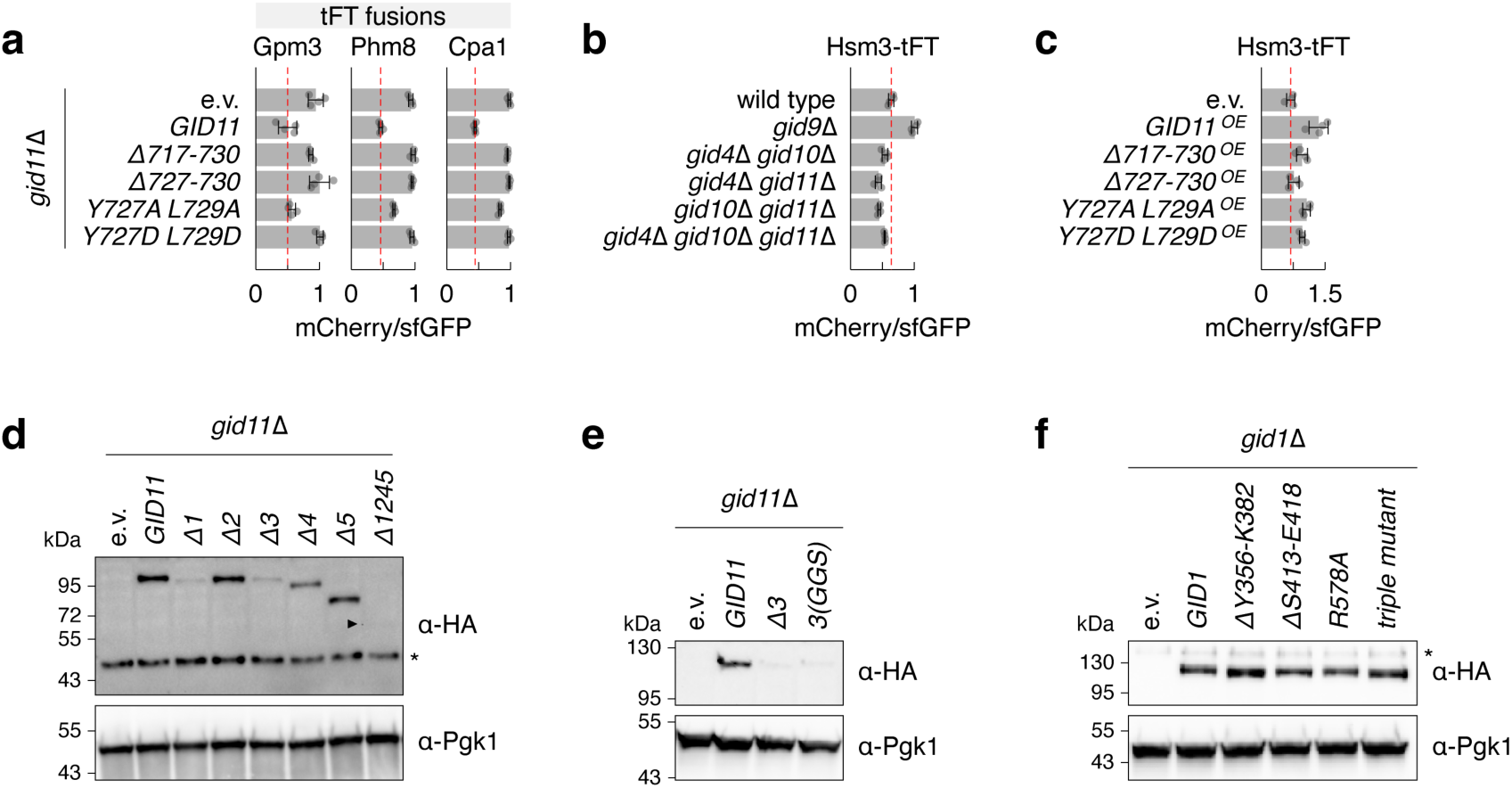
**a-c** – mCherry/sfGFP ratios of colonies expressing tFT-tagged proteins (mean ± s.d., n = 4 biological replicates; e.v., empty vector). OE, overexpression from the *GPD* promoter (**c**). **d-f** – Whole-cell extracts analyzed by immunoblotting 6 h after the glucose-to-ethanol switch (**d, e**) or from cells grown on glucose medium (**f**). Protein levels of HA-tagged Gid11 IDR mutants (**d, e**) or 3HA-tagged Gid1 mutants (**f**) (triple mutant – ΔY356-K382 ΔS413-E418 R578A). e.v., empty vector; OE, overexpression; *, non-specific background band; arrowhead, band corresponding to HA-Gid11-Δ1245.

**Figure S4.**
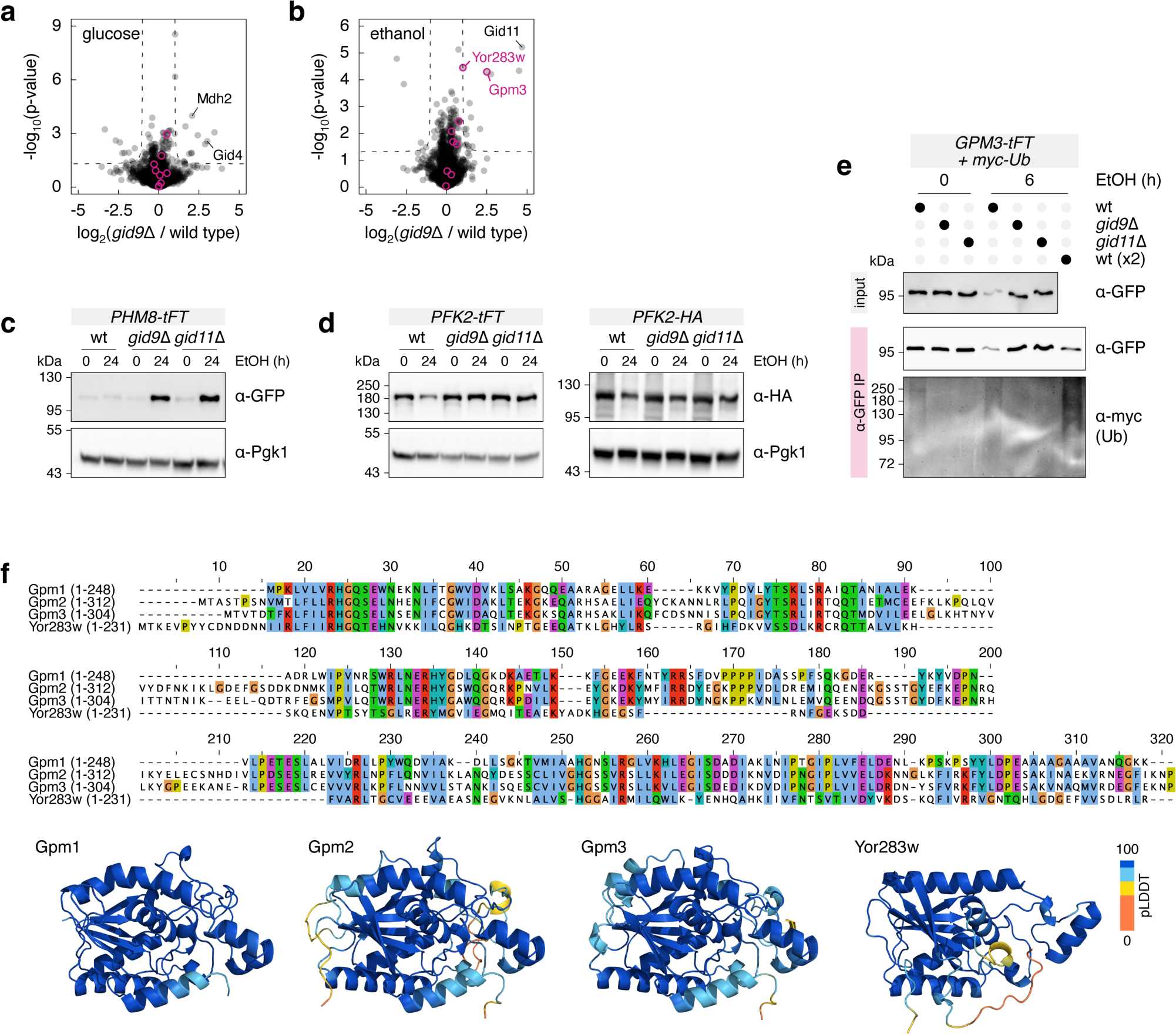
**a, b** – Differential proteomics of wild type and *gid9*Δ strains grown on glucose as carbon source (**a**) or 6 h after switching to ethanol (**b**). Gid11 substrates with a threonine N-terminus (Fig. 5a, with the exception of Phm8) are marked in magenta. **c, d** – Protein levels during the switch of carbon source from glucose to ethanol. Whole-cell extracts analyzed by immunoblotting. **e** – Ubiquitination of Gpm3 during the switch of carbon source from glucose to ethanol. Whole-cell extracts (input, top) and anti-GFP immunoprecipitates (IP, bottom) analyzed by immunoblotting. Myc-tagged ubiquitin was overexpressed in *GPM3-tFT* strains. **f** – Multiple sequence alignment (top) and AlphaFold 3 models (bottom) of *S. cerevisiae* Gpm1-3 and Yor283w.

**Figure S5.**
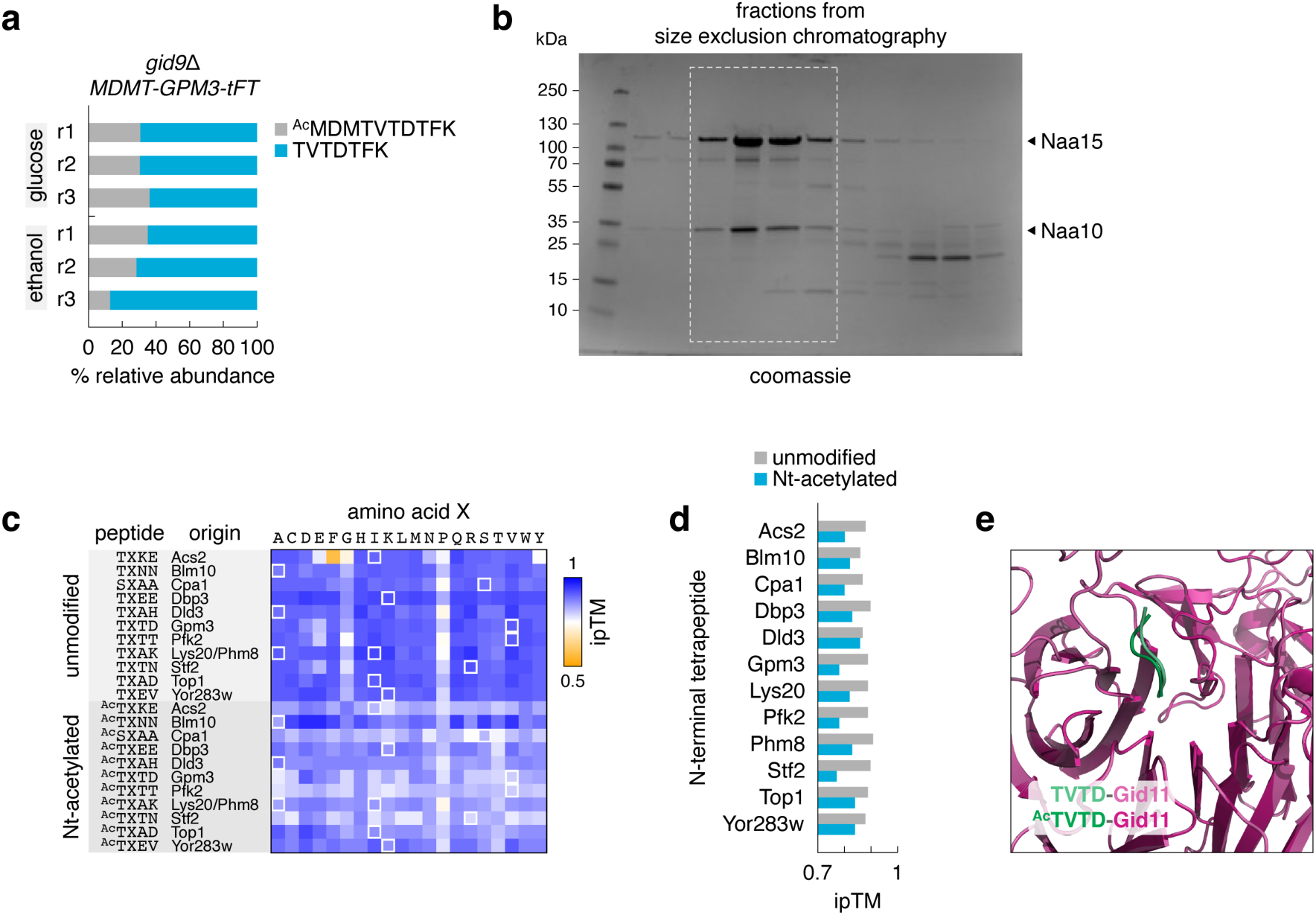
**a** – Mass spectrometry analysis of N-terminal peptides of the MDMT-Gpm3 variant. MTMT-Gpm3-tFT was immunoprecipitated from cells shifted from glucose to ethanol medium for 24 h. Percentage of spectral counts for the two detected N-terminal peptides (n = 3 technical replicates r1, r2 and r3). **b** – Purification of the Naa10-Naa15 complex. Fractions marked with a dashed outline were pooled and used in Nt-acetylation assays (Fig. 6e). **c, d** – Interface predicted template modeling (ipTM) scores of AlphaFold 3 models. Gid11 was modeled with tetrapeptides derived from N-termini of the indicated proteins. Peptide variants differing in the second amino acid, with or without an N-terminal acetyl group, were modeled with Gid11. Wild type second amino acids in the proteins of origin are highlighted with light outlines on the heatmap. ipTM scores of Gid11 models with the wild type tetrapeptides (**d**). **e** – AlphaFold 3 models of Gid11-Δ1245 with TVTD peptides, unmodified or Nt-acetylated, used as a starting point in molecular dynamics simulations.

**Figure S6.**
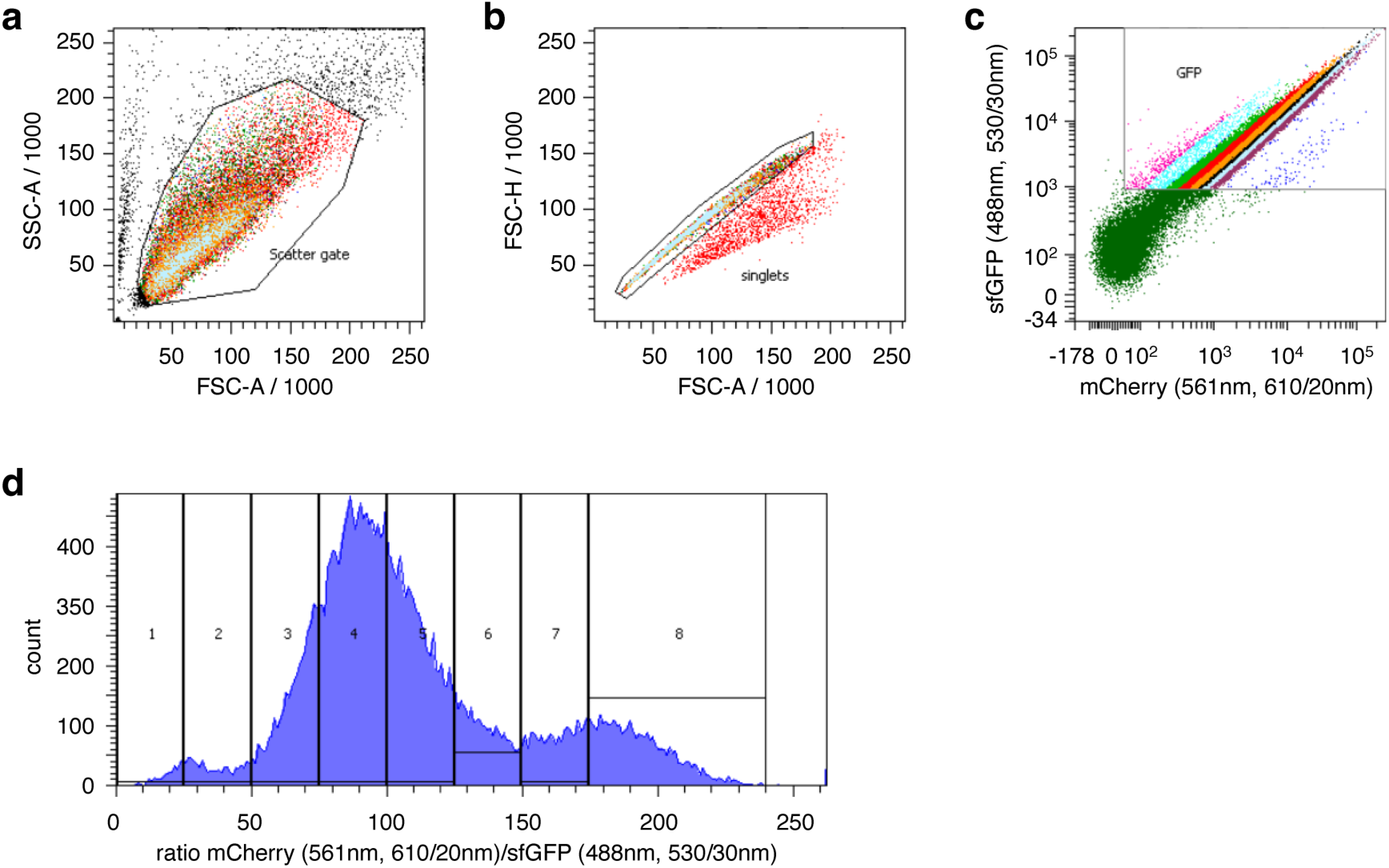
Representative gating strategy used for fluorescence-activated cell sorting in MPS profiling experiments, exemplified with the saturation mutagenesis library of the Gpm3 N-terminus. Gated cells (**a**) were subsequently gated for single cells (**b**), followed by gating for GFP-fluorescent cells (**c**). The resulting population was sorted into 8 bins according to the mCherry/sfGFP ratio (**d**). The 8 bins are also indicated with different colors in **c**.

## Supplementary Tables

Supplementary Table 1 – Yeast strains used in this work.

Supplementary Table 2 – Plasmids used in this work.

Supplementary Table 3 – Oligonucleotides used in this work.

## Supplementary Data

File Name: Supplementary Data 1

Description: SGA screens for regulators of GID11 expression; log_2_ fold changes in mCherry and sfGFP intensities, and corresponding p-values, between each mutant and the screen median.

File Name: Supplementary Data 2

Description: Composition of the GID^Gid11^ complex; mass spectrometry of Gid1-HA and HA-Gid11 immunoprecipitates; log_2_ fold changes in LFQ intensities, and corresponding p-values, between each sample and an untagged control immunoprecipitate.

File Name: Supplementary Data 3

Description: Whole cell proteomics with mass spectrometry of *gid9*Δ and *gid11*Δ mutants; log_2_ fold changes in LFQ intensities, and corresponding p-values, between each mutant and a wild type control.

File Name: Supplementary Data 4

Description: MPS profiling of Phm8 and Gpm3 N-terminal variant libraries in wild type, *gid9*Δ and *gid11*Δ backgrounds; N-terminal sequences of Phm8 and Gpm3 variants and corresponding PSIs.

File Name: Supplementary Data 5

Description: Archive file containing five text files in the GROMACS format MDP (molecular dynamics parameters) with the parameters used in the molecular dynamics simulations for the energy minimization (file name: emin_charmm36), NVT equilibration (file name: nvt_charmm36), NPT equilibration (file name: npt_charmm36) and production with (file name: production_vel_charmm36) and without (file name: production_novel_charmm36) velocity generation.

